# Regulation of epithelial tissue homeostasis by active trans-epithelial transport

**DOI:** 10.1101/2025.09.17.676587

**Authors:** Huiqiong Wu, Charlie Duclut, Gregory Arkowitz, Ranjith Chilupuri, Tien Dang, Jacques Prost, Benoit Ladoux, René-Marc Mège

## Abstract

Epithelia are intricate tissues whose function is intimately linked to mechanics. While mechanobiology has primarily focused on factors such as cell-generated contractility and mechanical properties of extracellular matrix (ECM), a novel mechanobiological paradigm highlights the role of osmotic and mechanical pressures in shaping epithelial tissues. In our study, we developed an in vitro model of cell coated micro-sized hydrogel spheres (MHSs) which allows to decipher the interplay between cellular activities and tissue mechanics. Drastic, isotropic MHS compressions were observed once the epithelia reached confluence. Further studies revealed that the compression was a process independent of cell contractility but rather regulated by active transepithelial fluid flow. Compressive stresses of about 7 kPa are generated by such an active hydraulic mechanism. Tissue homeostasis is then maintained by a fine balance between cell proliferation and extrusion. Our findings demonstrate the critical role of fluid transport in generating mechanical forces within epithelial tissues. Supported by a theoretical mechano-hydraulic model, a mechanistic framework for understanding the intricate interplay between cellular processes and tissue mechanics was established. These results challenge traditional views of epithelial tissue mechanics, emphasizing the pivotal influence of osmotic and mechanical pressures in shaping tissues. We anticipate that this study will advance the understanding of epithelial tissue development, the maintenance of homeostasis, and the mechanisms underlying pathological conditions.

**Significance Statement:** Epithelial tissues are vital for many bodily functions, but their mechanics remain poorly understood. Our study uncovers a novel mechanism by which epithelial cells generate mechanical stress, not through traditional cell contractility, but by actively pumping ions and water across their membranes. Using a model of micro-sized hydrogel spheres (MHSs) coated with epithelial cells, we demonstrate that epithelial cells actively transport fluid across the tissue to compress the MHSs. This facilitates the establishment of tissue homeostasis, which is further maintained by a balanced cell proliferation and extrusion rate. Supported by a theoretical model, our findings highlight the overlooked role of fluid transport in tissue mechanics, offering new insights into how epithelial tissues develop, maintain stability, and respond to disease.

## Introduction

Epithelial tissues serve as the protective linings of organs, creating vital barriers and functional interfaces between internal and external environments. Dysfunctions within epithelial layers lead to severe organ failures, making the understanding of their development and maintenance critically important. Apart from biochemical signaling, abundant literature points out the importance of mechanics on epithelial morphogenesis and homeostasis (1, 2). Physical properties of extracellular matrix (ECM) and neighboring tissues have been shown to regulate collective cell migration (3-5), tissue organization, epithelia folding (4-6) and integrity (7, 8).

Cells as active systems generate and adapt internal stresses and traction forces (9) in response to these mechanical cues. The underlying mechanisms rely on actomyosin generated forces (1, 2, 10, 11), as well as cell-matrix and cell-cell adhesions mechano-transduction and -sensing (12-14). However, cell contractility fails to explain some tissue behaviors highlighting alternative mechanisms, such as active ion and water transport through which cells can generate stresses (15-19). Despite the existence of fluid and osmolyte gradients in organs and tissues, including epithelia, the role of active ionic transport and water flow has been largely overlooked to understand epithelial cell behavior.

Fluid transport and mechanical pressure have been shown to be crucial for organ shaping including lung, kidney, vasculature and mammary glands (17, 20, 21). During murine lung development, the transmural pressure created by active fluid secretion of epithelial cells controls the airway branching (17). In regenerating *Hydra* spheroid, the inflation driven by transepithelial pumping creates mechanical stimuli that activate the Wnt3 signaling of head organizers (18). In the nematode germline, hydraulic instabilities among germ cells determine the cell fate (22). In developmental processes, such as inner ear and larva-polyp morphogenesis, epithelial tissues deposit highly charged ECM which creates osmotic pressures thereby promoting tissue morphogenesis (23, 24). Transepithelial fluid transport, generating hydraulic pressure within the inner cell mass, is a prerequisite for mammalian embryonic development (19, 25). Stress balance between the mechanical pressure in the lumen and tissue tension results in cyclic inflation-deflations, thereby regulating tissue and organ size (26, 27).

Epithelial cell lines are classical *in vitro* models to study fluid transport and lumen formation. In particular, MDCK (Madin-Darby Canine Kidney) cells are apicobasal polarized and behave as semipermeable membranes (28, 29). Their efficiency for fluid transport depends on a high transepithelial resistance, regulated by tight junctions, enabling the formation of transepithelial osmotic gradients (30, 31). In 3-dimensional (3D) environments, MDCK cells self-organize into cysts with polarities dependent on culture conditions, i.e. when cultured in ECM gels, their apical side faces the lumen while in suspensions their basal side faces the lumen (32). Regardless the polarity orientation, the lumen expansion relies on water influx that arises from ion and hydrostatic pressure gradients (33, 34). When cultured on impermeable 2D surfaces, MDCK cells form blisters or domes that result from cellular detachment from the underlying surface (35, 36). The formation of these blisters is caused by an increase of the basal hydrostatic pressure established by apical-to-basal fluid flux. This basal hydrostatic pressure nucleates local intercellular fractures, thus promoting the assembly of suspended multicellular structures that can sustain large deformations (37).

To further investigate the mechanical constraints imposed by both cellular contractility and hydraulic stresses on 3D epithelial tissues, we developed a novel synthetic model of 3D MDCK epithelial layers covering biofunctionalized MHSs (Micro-sized Hydrogel Spheres). We show that at confluency, epithelial cells actively deform the MHSs not through cellular contractility but rather through active pumping, leading to the build-up of an osmotic pressure difference that drives the deflation of the MHS. This compression relies on the establishment of an appropriate barrier function as well as ion and water transport across the epithelial tissue. After compression, the epithelium adapts to the reduced surface area not only by altering cell shape but also by maintaining a balance between cell division and extrusion as described in 2D cell cultures (38, 39). Our experimental results are supported by a theoretical model that incorporates both tissue tension and fluid transport. This model provides new insights into the mechanisms underlying both the maintenance of the homeostatic state and the observed fluctuations around it. Together, our results highlight the significant role of fluid transport in epithelial mechanics and consequent tissue homeostasis regulation.

## Results

### 3D epithelial monolayers compress MHSs via stresses independent of cell contractility

We first prepared fibronectin-functionalized polyethylene glycol diacrylate (PEGDA) MHSs by a simple emulsion synthesis (See Material and Methods). These MHSs were then sowed with epithelial cells and embedded in agarose to allow for live imaging (**Fig. S1**). Epithelial cells, initially sparsely distributed, spread well and proliferate on the MHSs up to confluency (**Fig. 1A, Video S1**). We observed that the MHSs systematically underwent substantial isotropic compression that only started after cells reached confluency (**Fig. 1B)**. Quantitative analysis demonstrated a consistent maximum reduction in MHS diameter of 21 ± 3% for MDCK cells and 23 ± 3% for human colorectal adenocarcinoma (Caco-2) cells (Fig. 1B, C, F; **Video S2**), irrespective of the initial size. These reductions correspond to volume decreases of 51% and 68%, respectively. Although the precise moment when cells reached confluency on each MHS could not be determined, the duration of compression—defined as the time from the onset of deformation to when the MHSs reached their minimum diameters—was approximately 6 hours. By 10 hours, ∼83% of the maximum deformation had been achieved (**Fig. 1D**).

**Figure 1.**
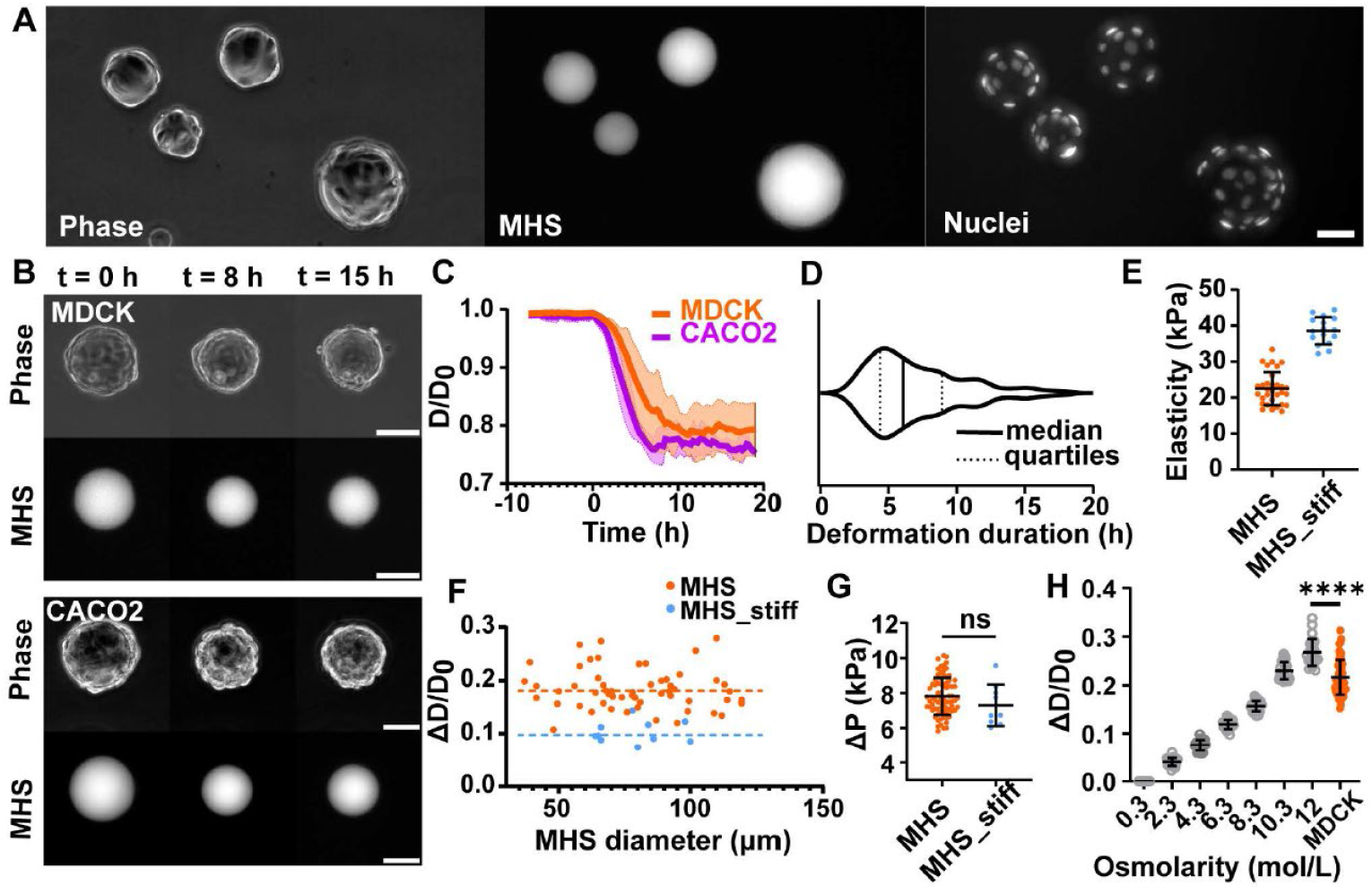
MHS deformation mediated by epithelial tissues. MDCK **(A)** and Caco2 **(B)**epithelial cells cultured on MHSs. Phase image (left), MHS channel (middle) and Nuclei channel (right); Scale bar: 50 µm. **(C)** Averaged evolution curves of the MHS diameters when cultured with MDCK (Orange, n = 12, N=3) or Caco2 cells (purple, n = 6, N = 1). Time is normalized to the onset of MHS size reduction. **(D)** Violin plots of deformation duration distribution with MDCK epithelia, n=123, N= 4. **(E)** Indentation measurement results of the Young’s modulus of two different MHS compositions, (MHS) used throughout the study when not specified n = 31, N = 3, MHS _Stiff) n = 16, N = 3. **(F)** MHS Deformations as a function of initial MHS sizes; orange dots represent MHSs with lower rigidity (n = 82, N =3) and blue dots represent MHS_stiff (n = 9, N = 1); The straight dotted-lines are the mean values, respectively. **(G)** Compressive stresses inferred from MHS deformations, n (MHS) = 58, N = 3; n (MHS_stiff) = 10; N = 1. **(H)** Deformations of bare MHS in deswelling experiments. Varied osmolarities were obtained by adding extra NaCl to the culture media; (MDCK): MHSs seeded with epithelial cells in media with physiological osmolarity (0.3 mol/L). n = 24, N = 4 for each condition without cells; n = 56, N = 3 for deformation with MDCK cells. ns: t test results, p>0.05; ****: t test results, P ≤ 0.0001.

We then calculated the amount of stress developed by the tissue to compress the MHSs. We determined the Young’s modulus of the MHSs to be 22.5 ± 4.4 kPa using nano-indentation **(Fig. 1E)**. Knowing the elastic modulus, we inferred from the observed maximal compression the compressive stress to be 7.8 ± 1 kPa (**Fig. 1G)**. To investigate whether this mechanical deformation was dependent on hydrogel stiffness, we prepared stiffer MHSs_stiff (Young modulus = 38.6 ± 3.6 kPa) **(Fig. 1E)**. When MDCK cells reached confluency on these stiffer substrates, we observed a maximum reduction in MHS diameter of 10 ± 2% regardless of the initial MHS size (**Fig. 1F**), corresponding to a tissue-generated compressive stress of 7.3 ± 1 kPa (**Fig. 1G**). These results show that tissue-generated stresses are independent of substrate stiffness. Overall, our findings suggest that the stress driving MHS deformation may be independent of cell contractile-based forces, which have been reported to be rigidity-dependent at the single cell (40-42) and collective cell levels (3, 14). Next, we explored whether the final compression state of MHSs was determined by the material properties of the hydrogel or governed by the epithelial tissue covering it. Simple deswelling experiments (43) revealed a progressive reduction in MHS diameters when naked MHSs were incubated in media with increasing NaCl concentrations (**Fig. 1H**). Notably, to obtain significant radius reductions, as those observed in the presence of an epithelium, one needs to reach extremely high osmolarities, typically 10 mol/L ruling out a direct effect of deswelling on the hydrogel.

Next, we delved into understanding the underlying source of the stress responsible for the deformation of the MHSs. To further investigate the role of actomyosin-based contractility, we inhibited non-muscle myosin II (NMM II) activity using blebbistatin, applied either before or after compression. This treatment had no significant effect on the maximal compression achieved (**Fig. 2A, D**). Similarly, treatment with calyculin A, a phosphatase inhibitor that enhances cellular contractility, did not affect MHS compression (**Fig. 2B, D**). These findings reinforce the hypothesis that acto-myosin contractility is not the driving stress behind MHS deformation. Finally, we observed that MDCK cells knockdown for NMM IIA (NMM IIA KD) (44) compressed the MHSs as efficiently as WT cells (**Fig. 2C-D**). Altogether, these results demonstrate that the MHS compression could not result from contractile forces generated by the actomyosin cytoskeleton.

**Figure 2.**
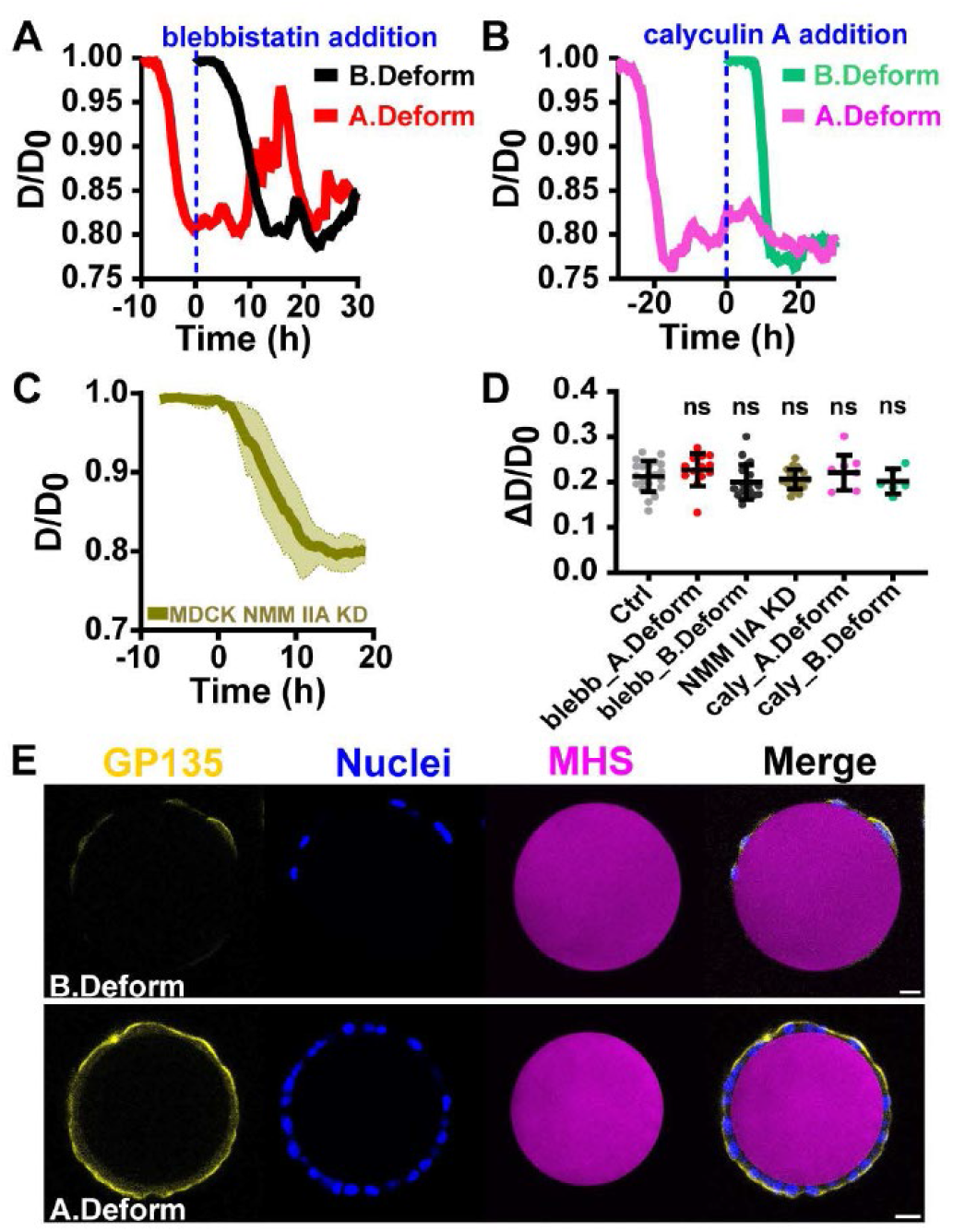
Tissue contractility modulation. **(A)** Representative size evolution curves of MHSs under blebbistatin treatment (20 µM), drug was administrated at t = 0 h. **(B)** Representative size evolution curves of MHSs under calyculin A treatment (5 nM), drug was administrated at t = 0h. **(C)** Averaged size evolution curves of MHSs seeded with MDCK NMM IIA KD cells, here time is normalized to the start of each deformation, N = 2, n = 9. **(D)** Quantification of deformations under different contractility modulation conditions, (A.Deform) n = 13, N=3; (B.Deform) n = 19, N=3; (NMM IIA KD) n = 23, N = 2; (caly.A.Deform) n = 9, N = 1; (B.Deform) n = 6, N = 1. B.Deform and A.deform stands for administration before and after deformation respectively. **(E)** Immunostaining of apical marker GP135, before deformation (B.Deform) and after deformation (A.Deform). Scale bars = 20 µm.

### Proper barrier function, osmotic gradient and water outflow are required for MHS compression

To induce compression of the MHSs, regardless of the source of the mechanical stress involved, the incompressible fluid must be expelled from the MHSs. High-resolution confocal live imaging of MDCK LifeAct-GFP (**Fig.S2**) revealed that MHS compression coincides with a reduction in MHS-tissue total volume, indicative of a transepithelial fluid outflow (**Fig. 3A**). MDCK cells possess active transepithelial ion transport abilities (28, 29, 36). From MHS compression rate, we estimated an average water efflux rate per unit area of 0.27 ± 0.13 µL·h^−1^·cm^−2^, consistent with previously reported flux rates for MDCK cysts in 3D cultures (0.25 ± 0.03 µL·h^−1^·cm^−2^ (45), 0.22 ± 0.01 µL·h^−1^·cm^−2^ (46)). Reported flux rates for 2D MDCK monolayers vary widely from 0 to 10 µL·h^−1^·cm^−2^ (47), depending on culture conditions and measurement methods, although higher values can be obtained in a apical-basal zero-pressure difference condition (48). We therefore hypothesized that active electrolytes transport across the cell monolayer establishes a transepithelial osmotic gradient, resulting in subsequent water outflow leading to MHS compression. If MHS compression is due to transepithelial outflow governed by osmotic gradients, the maximal compression of MHSs should change upon sudden changes in external osmolarity. When applying a series of hypotonic and hypertonic shocks, we indeed observed significant changes in MHS compression **(Fig. 3B, S3)**. Hypertonic shocks at 0.45 and 0.4 mol/L caused further MHS compression compared to physiological osmolarity (0.3 mol/L). Conversely, hypotonic shocks led to MHS relaxations, decreasing the equilibrium MHS compression to 19.0 ± 3.1%, 13.6 ± 2.1%, and 11.7 ± 1.3% for 0.2, 0.15, and 0.1 mol/L, respectively.

**Figure 3.**
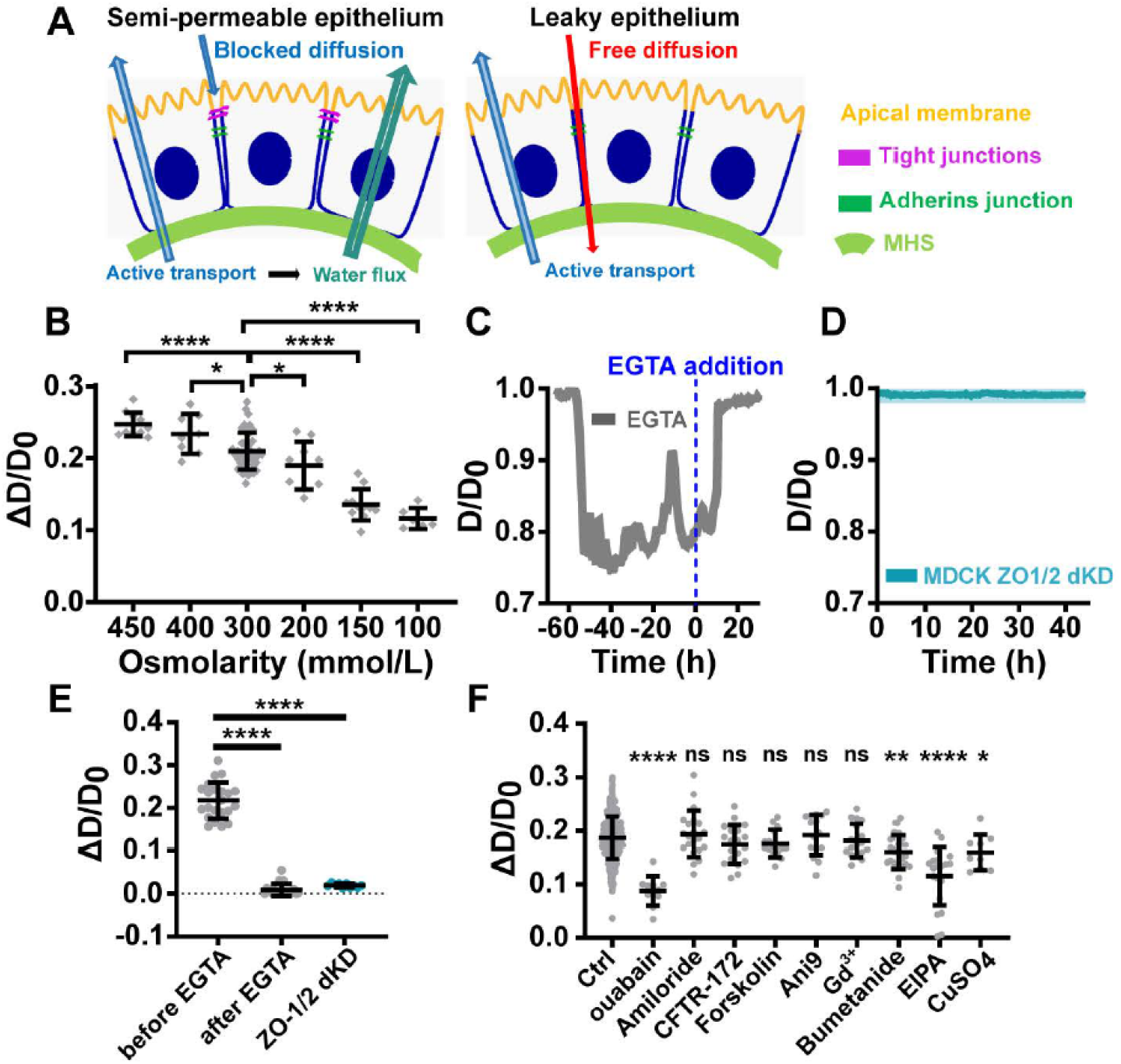
Tissue barrier function and active transport in regulating MHS deformations. **(A)** Schematics of semi-permeable and leaky epithelia and corresponding substance transports. **(B)** Quantification of deformations under different osmotic shock conditions. n (0.45 mol/L) =11, N =1; n (0.4 mol/L) = 8, N = 1; n (0.3 mol/L) = 45, N = 3; n (0.2 mol/L) = 9, N = 1; n (0.15 mol/L) = 11, N = 1; n (0.1 mol/L) = 6, N = 1. **(C)** Representative diameter evolution curve of MHS when treated by EGTA (3 mM). **(D)** Averaging diameter evolution curves over time of MHSs seeded with MDCK ZO-1/2 dKD cells. n = 10, N = 2. **(E)** The quantification of deformation under different barrier modulation conditions. n (EGTA) = 22, N = 2; n (ZO1/2 dKD) = 10, N = 1. **(F)** Quantification of deformation under different ion channel inhibition conditions, all the inhibitors were administrated before MHS deformations except for ouabain. ouabain) n = 11, N = 1; Amiloride n =21, N = 2; CFTR-172 n= 19, N= 2; Forskolin n = 15, N = 2; Ani9 n = 15, N = 2; Gd^3+^ n = 17, N = 2; Bumetanide n = 22, N = 2; EIPA n = 20, N =1; CuSO_4_ n = 9, N = 2.

To establish a transepithelial osmotic gradient, cells first need to acquire polarized transport ability which is linked to their apical-basal polarity (**Fig. 3A**). Immunostaining for the apical marker podocalyxin (GP135) (49) revealed that the cells were polarized, with the GP135-positive apical side oriented toward the culture medium, even before the onset of MHS compression (**Fig. 2E**). Non-confluent MDCK cells do not have the ability to build transepithelial osmotic gradients due to para-cellular free diffusion. For the establishment of such gradients, the epithelial monolayer needs to establish proper barrier function associated to the establishment of an uninterrupted tight junction belt (28, 30). Immunostaining of ZO1 indeed revealed the formation of apical tight junctions in the epithelial cell monolayer (**Fig. S4**). Furthermore, the perturbation of the barrier function by EGTA (3 mM), which destabilizes adherens and tight junctions (50), induced a complete relaxation of MHSs to their initial sizes (**Fig. 3C,E, S5A**). In line with this observation, we conducted experiments using MDCK ZO-1/2 double knockdown (dKD) cells (51), which lack functional tight junctions. These cells showed no MHS compressing ability, demonstrating the necessity of an intact epithelial barrier for MHS compression to occur (**Fig. 3D,E, S5B**). Overall, our findings reveal that MHS compression is driven by an osmotic gradient and active water outflow generated by a polarized epithelial monolayer, functioning as a semipermeable membrane.

### Active transepithelial transport drives MHS compression

To better understand the active and passive polarized transport across the epithelial monolayer, we performed immunostaining on compressed MHS (**Fig. S6**). The sodium-potassium pump Na^+^/K^+^-ATPase (NKA), which establishes essential electrochemical gradients for other ion channels (52), displayed a distinct basolateral distribution. In contrast, the sodium-potassium-chloride cotransporter 1 (NKCC1), a key regulator of ion homeostasis in mammalian tissues (53), and the sodium-hydrogen exchanger 1 (NHE1), an important cell volume and pH regulator (54), were predominantly localized on the apical side.

To confirm that active transport across the epithelial monolayer drives MHS compression, we pharmacologically inhibited various ionic pumps and channels. Treatment with ouabain, a specific inhibitor of Na+/K+-ATPase (NKA), administered after MHS compression, resulted in significant MHS relaxation (**Fig. 3F, S7A**). Conversely, amiloride, an inhibitor of the epithelial sodium channel (ENaC), primarily involved in sodium reabsorption in kidney and lung epithelia, had no significant effect on MHS compression (**Fig. 3F**). Similarly, neither inhibition by CFTR-172 nor activation by forskolin of the chloride channel CFTR (cystic fibrosis transmembrane conductance regulator), nor inhibition of another chloride channel, TMEM16A (55), by Ani9, nor inhibition of mechanosensitive ion channels by Gd^3+^ (38), affected MHS compression (**Fig. 3F**).

In contrast, inhibition of the sodium-potassium-chloride cotransporter NKCC1 with bumetanide significantly reduced MHS compression to 16.0 ± 3.1 % compared to 18.7 ± 4.0 % for controls, highlighting its critical role in the deformation process **(Fig. 3F, S7B)**. Similarly, inhibiting the sodium-hydrogen exchanger 1 (NHE1) with EIPA led to a pronounced reduction in MHS compression (11.5 ± 5.3 % compared to 18.7 ± 4.0 % for controls) (**Fig. 3F, S7B**). According to our hypothesis, transepithelial ion transport should drive passive water outflow facilitated by water channels such as aquaporins. Consistent with this, immunostaining showed aquaporin 3 (AQP3) localized in both basolateral and apical domains (**Fig. S6**). Inhibition of AQP3 using CuSO4 (56) caused a mild but significant reduction in MHS compression (15.9 ± 3.2% compared to control conditions 18.7 ± 4.0 %) (**Fig. 3F, S7B**). Altogether, these findings identify key ionic transporters and pumps involved in MHS compression. They provide compelling evidence that MHS compression results from active transepithelial ion transport, which generates osmotic and mechanical pressures essential for this process.

### Tissue maintains long term homeostasis after reaching mechanical steady state

The decrease in MHS size resulted in a reduction in the epithelial monolayer surface area, averaging 37%. To investigate how this self-generated compression affects the epithelium organization and dynamics, we analyzed changes in cellular morphology before and after compression. Following MHS compression, we observed a major reduction in apical cell area (214.0 ± 105.1 µm^2^ at compressed state vs. 345.0 ± 215.5 µm^2^ at the onset of compression), accompanied by an increase in cell height (10.8 ± 3.4 µm at compressed state vs. 6.8 ± 2.9 µm at the onset of compression), indicative of a transition to a more columnar morphology (57, 58) (**Fig. 4A-C**). Moreover, cell volume estimation computed from cell surface area and height revealed no statistically significant changes upon MHS deformation. Live imaging further revealed that cell height started to increase before confluency (**Fig. S8**), indicative of a progressive maturation of the epithelium. During compression, cell morphological changes were purely geometrical since they occurred at constant cell volume. To further investigate the impacts of MHS compression on the epithelium, we measured cell densities at various time points: the onset of compression, the end of compression, and 2.5, 5- and 10-hours post-compression (**Fig. 4D**). Consistent with the reduction in apical cell area, cell density nearly doubled immediately after compression (43.5 ± 4.8 cell/10^4^ µm^2^) compared to the onset (24.8 ± 4.8 cell/10^4^ µm^2^). The density then remained stable for at least three days **(Fig. 4D, Fig. S9A)**, similar to that observed on 2D PAAm surfaces (**Fig. S9B**). Interestingly, cell densities at both the onset and end of compression were independent of the initial MHS sizes, indicative an intrinsic tissue property (**Fig. 4E**). Comparable density evolutions of Caco2 epithelia on MHSs and on 2D substrates were also observed **(Fig. S9C-D)**. Additionally, live observations revealed a remarkable decrease in cell motility, from an average cell velocity of 14.6 ± 7.8 µm/h before compression to 4.2 ± 3.5 µm/h afterward (**Fig. 4F**). The concomitant decrease in cell motility and increase in cell density suggest that the 3D monolayers on MHSs transitioned to a homeostatic state.

**Figure 4.**
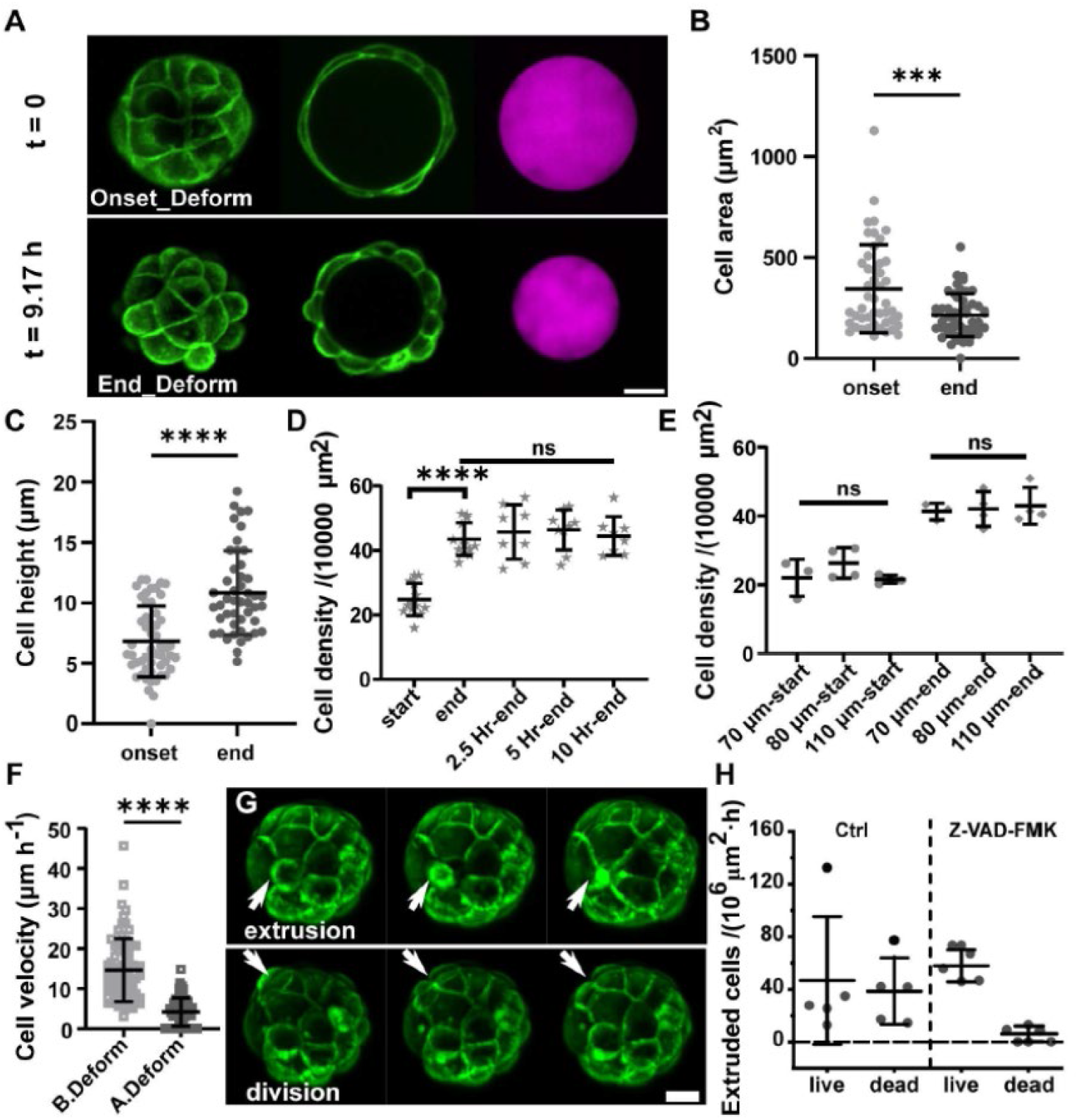
Tissue homeostasis on the MHSs. **(A)** Representative evolution of MDCK CAAx-GFP cell morphology changes on MHS at the onset (t=0 min) and the end (t=550 min) of deformation, from left to right it shows the hemi-MHS z projection of cells, cells on the equatorial plane of the MHS, and corresponding MHS channel, respectively. **(B)** Cell area quantification at the onset and end of deformation. n(onset) = 47, N=4; n (end) = 51, N=4. **(C)** Cell height quantification at the onset and end of deformation. n(onset) = 52, N = 4; n(end) = 47, N=4. **(D)** Cell density at the onset of deformation, the end of deformation and 2.5, 5, 10 hours -post deformation, n(onset) = 13, N = 2; n(end) = 13, N = 2; n(2.5 Hr-end) = 8, N = 2; n(5 Hr-end) = 9, N = 2; n(10 Hr-end) = 8, N = 2. **(E)** Cell density on MHS with varied initial sizes at the onset of deformation and the end of deformation. **(F)** Cell velocity quantification before and after deformation. n(B.Deform) = 78, N = 2; n(A.Deform) = 61, N = 2. **(G)** Representative cell extrusion and division events after MHS deformation. **(H)** Quantification of extruded cell fate expressed as frequency per unit area under control conditions and in Z-VAD-FMK treated conditions, n(ctrl) = 5, N = 1; n(Z-VAD-FMK) = 5, N = 1. Scale bars = 20 µm.

Tissue crowding has been reported to induce cell extrusion (38, 59, 60). Performing live imaging, we observed a surge of cell extrusion events following MHS compression (**Fig. 4G, Fig. S10, Video S3**). To investigate the fate of extruded cells, we labeled them with annexin V antibodies (60). Both live and dead cell extrusions were observed (**Fig. 4H**). Tissue homeostasis, as revealed by cell density maintenance over 3 days post MHS compression (**Fig. 4D, Fig. S9**), implies a balance between cell extrusion and division. Counting these events revealed indeed a balance between cell extrusion rates (41 ± 16 cells/ (10^6^µm^2^·h)) and cell division rates (41 ± 18 cells/ (10^6^µm^2^·h)) on compressed MHSs (**Fig. S11, Video S4**). Interestingly, fluctuations in bead compression correlated with cell extrusion and division events (**Fig. S10, S11**), highlighting the dynamic nature of this self-regulated homeostatic state. To explore how the tissue adapts to perturbations, we inhibited apoptosis using Z-VAD-FMK (61). Apoptotic blockade induced a predominance of live-cell extrusions versus dead cell extrusions, with however no significant effect on maximum MHS compression (**Fig. 4H, Fig. S12**). These results suggest that the homeostatic control of the closed epithelial monolayer is a robust and adaptable process.

Our findings demonstrate that confluent epithelial tissue actively generates transepithelial fluid outflow, leading to the compression of the MHS it resides upon. This compression process is accompanied by tissue maintenance of close to *in vivo* homeostatic state, where cell division and extrusion are actively balanced.

### A spherically-symmetric model for tissue mechanics and hydraulic properties recapitulates the experimental data

To further determine the mechanisms at play, we developed a theoretical model that incorporates both tissue contractility and fluid transport. In this description, the cell height is neglected, and the epithelial cell monolayer is considered as a spherical semi-permeable membrane of radius *R*(*t*) enclosing a soft gel (MHS) (**Fig.5A**).

**Figure 5.**
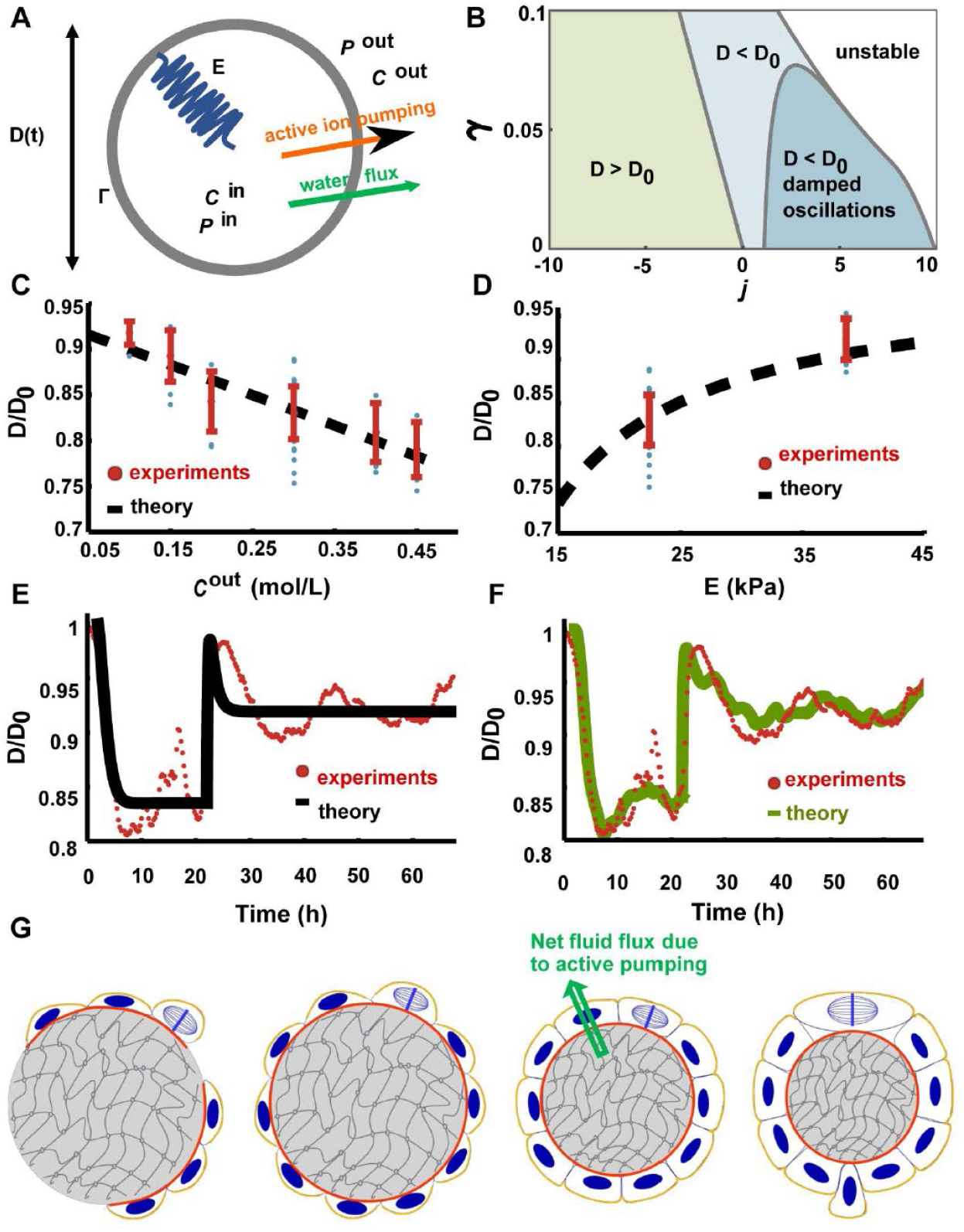
Mechano-hydraulic model of a tissue monolayer. **(A)** Schematics of the theoretical model. Cells actively pump ions from the basal to apical side (orange-color arrow), which leads to a water efflux (green arrow) following osmotic pressure difference. This water efflux results in gel compression thus generating a compressive stress, which is symbolized by a spring. **(B)** State diagram of the model for different values of the dimensionless pumping *j* = *σJ*^*p*^*C*^*out*^/3*B*Λ_*i*_ and surface tension *γ* = Γ/3*BR*_0_. **(C)** Normalized steady-state radius *D*/*D*_0_ as function of the medium concentration *C*^*out*^ (in mol/L). The black dashed line is a fit of Eq. (4) to the experimental data (dots). Fitted values: *J* = *σJ*^*p*^/ 3*B*Λ_*i*_ = 3.3 10^−1^ L/mol and *γ* = Γ/3*BR*_0_ = 3.8 10^−2^(dimensionless). **(D)** Normalized steady-state radius *D*/*D*_0_ as function of the gel Young’s modulus *E* (in kPa). The black dashed line is a prediction from Eq. using the values fitted in (C). **(E)** Normalized radius *R*(*t*)/*R*_0_ as function of time during an osmotic shock experiment (red dots). At *t* = *t*_*shock*_ ≈ 22*h*, the medium is diluted from *C*^*out*^ = 0.3 mol/L to *C*^*out*^ = 0.1 mol/L. The black line shows the corresponding dynamics obtained from fitting the theoretical model. **(F)** Normalized radius *D*(*t*)/*D*_0_ as function of time during an osmotic shock experiment (red dots, same data as in (E)). The green solid line shows a realization of the stochastic model (picked among 1000 realizations) that resembles the experimental data. **(G)** Schematics representing the whole self-regulated homeostasis process due to active fluid transport in epithelial tissue. See SI for details.

We first discuss the mechanical properties of the system. The isotropic compression of the MHS results in an elastic stress proportional to the relative deformation that reads 3*B*(*R*_0_ − *R*(*t*))/*R*_0_ where *B* is the gel bulk modulus and *R*_0_ = *R*(*t* = 0) is the radius of the MHS prior to compression. This isotropic stress must be balanced by the Laplace pressure contribution 2Γ/*R*(*t*) where Γ is the tissue surface tension stemming mainly from the acto-myosin cortex contractility, and by the mechanical pressure difference *P*^*in*^ − *P*^*out*^ between the inside and of the tissue layer. Force balance thus reads:

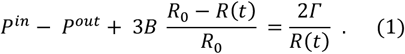

To account for active tissue pumping, an irreversible thermodynamics framework (62, 63) is used. Cells actively pump ions, which in turns creates an osmotic pressure difference leading to (passive) water flux. Water flux is thus driven by two thermodynamic forces: mechanical and osmotic pressure differences, such that the sphere volume *V*(*t*) = 4*πR*(*t*)^3^/3 dynamics obeys:

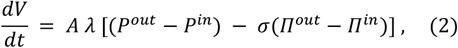

where *A* = 4*πR*(*t*)^2^ denotes the sphere surface area, *λ* is the tissue permeability to water flows, and Π^*out*^ − Π^*in*^ = − *k*_*B*_ *T ΔC* with *ΔC* = *C*^*in*^ − *C*^*out*^ denotes the osmotic pressure difference across the epithelium. Particularly, we have included a reflection/selectivity coefficient *σ* (63, 64). A fully semi-permeable membrane that allows only water to pass corresponds to *σ* = 1, while a fully permeable membrane that is permeable to both water and osmolytes has *σ* = 0. When accounting for ion diffusion, which may influence the generated osmotic pressure, we determined that ion concentrations within the MHS reach homogeneity on a timescale of approximately one second. By contrast, equilibration in the outer medium requires up to 42 hours. Incorporating diffusion into the model results in a slightly accelerated compression but does not substantially affect the overall conclusions (see SI Theoretical model). Accordingly, in the subsequent analysis, we assume that the concentrations C^in^ and C^out^ are homogeneous within the MHS and the outer medium, respectively.

Finally, an osmotic pressure difference builds up due to cell active pumping. For simplicity, we consider here the transport of a single ionic species. The number *N*^*in*^ of ions in the sphere evolves according to:

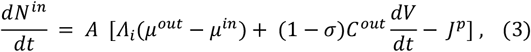

where Λ_*i*_ is the Onsager coefficient for ion transport through the membrane, *μ*^*out*^ − *μ*^*in*^ ≈ − *k*_*B*_*T ΔC*/*C*^*out*^ denotes the chemical potential difference between the outside and inside and *J*^*p*^ is the flux due to active transport (positive when directed outwards).

This system of equations captures sphere compression with a stable steady-state radius *R*^*\**^ < *R*_0_ when the active ion flux is directed outwards (*J*^*p*^ > 0), or sphere expansion when the active ion flux is directed inwards (*J*^*p*^ < 0) and surface tension is sufficiently low. **Fig. 5B** displays a state diagram of our model for different values of the parameters. In our experimental system, robust MHS compressions are observed, corresponding to the epithelial active transport being directed outwards (*J*^*p*^ > 0), increasing the concentration outside and therefore creating a nonvanishing osmotic pressure difference which leads to water efflux from the MHS. The normalized steady-state diameter can be obtained from our model and reads:

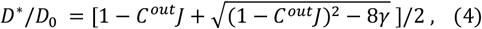

where *J* = *σJ*^*p*^/ 3*B*Λ_*i*_ and *γ* = Γ/3*BR*_0_. We then fitted Eq. (4) using experimental data for different osmotic conditions (**Fig. 5C**), thus obtaining *J* = 3.3 · 10^−1^ L/mol and *γ* = 3.8 · 10^−2^ (dimensionless) which are in agreement with literature values (62, 63). Note that we have considered a constant Onsager coefficient Λ_*i*_ rather than a constant membrane permeability to ion flows Λ = k_B_T Λ_*i*_ /*C*^*out*^. This choice yields the correct concentration-dependence of the steady-state diameter.

Without further fitting, Eq. (4) was then used to predict the steady-state radius for different MHS stiffnesses, and a remarkable agreement was shown with the experimental results (**Fig. 5D**). The fitted parameters from osmotic shock experiments **(Fig. 5C)** accurately capture the dependence of the steady-state radius on gel stiffness, especially for stiffer gels. However, additional experiments revealed a deviation at low stiffness (5.2 kPa), where tissues compressed less than expected **(Fig. S13E)**, suggesting that epithelial hydraulic properties themselves depend on the substrate stiffness, likely through reduced pumping or increased leakage.

We then compared the dynamics predicted by our model to the experimental ones. An example of this comparison is displayed on **Fig. 5E** for the case of an osmotic shock where the outer medium is diluted from *C*^*out*^ = 0.3 mol/L to *C*^*out*^ = 0.1 mol/L during the experiment. The fitting shows a good agreement with the first compression stage. Similarly, good fits are obtained for other osmotic shock experiments. Note however that the model accounts neither for the slower relaxation to the steady state that follows the osmotic shock, nor for the overall large fluctuations around the steady-state radius.

As shown in the experiments, the compression of MHS is coupled with cell density changes, and we thus further extended our model by including cell density fluctuations. The compression leads to an increase in cell density and enables a homeostatic state to be achieved rapidly. Therefore, close to the homeostatic state, we write the dynamics for the cell density *ϱ*(*t*) as a Langevin equation (see SI for details):

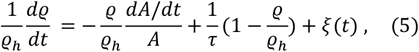

where *ϱ*_h_ is the homeostatic cell density. In Eq. (5), the first term on the right-hand side accounts for density variations due to MHS surface area changes, the second one for the relaxation to the homeostatic state with characteristic time *τ*, and the last term *ξ*(*t*) is a stochastic Gaussian white noise with vanishing mean and variance < *ξ*(*t*)*ξ*(*t′*) >= 2*W δ*(t − t*′*), where the noise amplitude *W* is expected to scale as 1/*N* where *N* is the number of cells. The feedback of cell density fluctuations to the mechano-hydraulic model then takes the form of a density-dependent surface tension:

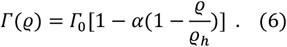

Where *α* is a dimensionless parameter and Γ_0_ is the surface tension at the homeostatic density. For *α* > 0, which we consider in the following, the tissue surface tension is lowered (Γ(*ϱ*) < Γ_0_) whenever the tissue density is smaller than its homeostatic density (*ϱ* < *ϱ*_h_).

An example of realization of the stochastic mechano-hydraulic model given by Eqs. (1-3) and (5-6) is provided in the SI. Large fluctuations around the mean steady-state radius, similar to those in the experiments, are observed. Those large fluctuations can be understood as a consequence of the positive feedback between density changes and surface tension: If a cell density fluctuation lower *ϱ*, it causes a lowering of the surface tension, which itself favors an increase of the radius and thus a further decrease of the cell density.

Finally, we return to the experimental data and compare our stochastic model to osmotic shock experiments. Notably, adding stochastic cell division/extrusion and their mechanical feedback leads to radius fluctuations that are similar to those observed experimentally (**Fig. 5F**). In addition, the stochastic model also accounts for the slower relaxation to the steady state that follows the osmotic shock. As an alternative to the density dependence of the surface tension in Eq. (6), we have considered other mechanisms by which density fluctuations could feedback on the hydraulic model. Since cell divisions or extrusions could trigger leakage (26, 27), we have also considered a density dependence of the Onsager coefficient *Λ*_*i*_ or of the permeability to water *λ*. In the SI, we show that a density-dependent Onsager coefficient *Λ*_*i*_ leads to similar results as a density-dependent surface tension. On the other hand, a density-dependent permeability to water flow only plays a role while the sphere volume is changing and thus does not generate radius fluctuations as observed in the experiments.

In summary, utilizing a simplified stochastic mechanical-hydraulic theoretical model, we managed to recapitulate the experimental observations, which validates our experimental conclusions. In particular, we note that the fitted dimensionless surface tension *γ* is indeed small and its contribution to the MHS compression is negligible compared to the role of ion pumping that leads to an osmotic pressure difference. This is also in agreement with the experimental observation that the Na+/K+-ATPase inhibition by ouabain leads to the most drastic reduction of the MHS compression: from the model perspective, inhibiting this pump means a smaller value of the parameter *J*^*p*^ in Eq. (3), which results in smaller steady-state diameter according to Eq. (4). In addition. the theoretical model provided an interesting perspective on the relationship between tissue surface tension and corresponding cell density, which can be useful in future studies.

## Discussion

Our study uncovers a hydraulic origin of stress generation in epithelial tissues. Using MHSs as a boundary-free model, we show that epithelia are capable of generating large isotropic compressive stresses, on the order of several kilo-Pascals, through active ion transport and subsequent water efflux. This finding challenges the long-standing view that acto-myosin contractility is the dominant driver of epithelial mechanics and suggests that hydraulic processes play a much larger role in tissue homeostasis than previously recognized.

Compared to prior reports on stresses generated by tissue hydraulics (26, 33, 48, 65), the stresses we measured here exceeds by more than an order of magnitude. In our system, the measured pressure reflects gel compression rather than hydrostatic pressure. Active basal-to-apical ion pumping generates an osmotic gradient that drives water efflux from the MHS, compressing the gel until osmotic and gel compression generated mechanical stresses equilibrate at ∼7 kPa. This compression also affects the MDCK monolayer, reinforcing columnar cell packing, tight junctions and barrier integrity, which explains the quasi-stable MHS size observed post-deformation. By contrast, in other systems such as domes and cysts, the accumulation of fluid stretches the cell monolayer, thus resulting in weakened cell-cell junctions that lead to junctional fracture which prevent higher pressure build-up, which could explain the observed cycle of inflation/deflation. In 2D systems, the previously reported lower pressures may result from fluid leaks that prevent osmotic gradients from building up. In addition, in cysts, underestimation could also be methodological: pipette aspiration measures local cortical tension rather than osmotic pressure, while efflux-based assays after puncture can underestimate pressure if the surrounding medium is porous and dissipates flow.

Our results also highlight a striking functional outcome: self-generated compression drives epithelia to a defined homeostatic state. The active ion transport and water efflux led to a self-compression of the monolayer by ∼37%, resulting in a homeostatic cell density that remained stable for at least 3 days, indicating that tissue self-compression achieved a transition to a preferred homeostatic density. Caco2 epithelia behaving similarly in maintaining cell density post MHS compression, suggests that active fluid transport may serve as a general regulator of epithelial homeostasis across tissues. Notably, this cell density was independent of MHS size, indicating that it was an intrinsic tissue property. Unlike externally applied compression, which induces transient density changes and often destabilizes tissues (38), transport-mediated self-compression produced a stable balance of proliferation and extrusion. Both live and dead cell extrusion were observed at higher rates than in 2D cultures (59), with a greater proportion of live cell extrusions (60). The observed increase in live-cell extrusion when apoptotic extrusion was inhibited without effect on self-compression further indicates that homeostasis is regulated independently of specific extrusion mechanisms.

This rapid establishment of homeostasis could not be achieved by proliferation alone, but was facilitated by active basal-to-apical fluid transport (**Fig. 5G**). Indeed, during self-compression, the epithelium reached a two-fold increase in cell density in 6 hours on average which is at least 3 times faster than would be possible through proliferation alone, which typically requires 18–24 hours (47). A particularly interesting aspect of this process is the directionality of fluid transport. MDCK epithelial tissues are known to transport fluid in both directions: basal to apical and apical to basal, depending on culture conditions. In conventional 2D cultures, whether on impermeable substrates (such as plastic, glass and PDMS) (65) or permeable supports (such as Transwell) (48), MDCK monolayers typically show net apical to basal transport. In contrast, 3D culture systems often display bidirectional transport. For example, MDCK cysts in suspension cultures generally transport from apical to basal, resulting apical-out cysts, whereas ECM-embedded cysts exhibit basal to apical transport, forming basal-out cysts (32). The factors that determine transport direction in these systems remain unclear, and further studies are needed to elucidate the underlying mechanisms.

To address the question of how fluid transport couples to mechanics at the tissue level, our modeling shows that even a small total trans-epithelial ion concentration difference (*C*^*in*^ − *C*^*out*^ ≃ −3 *m*M) is sufficient to drive water efflux and counter balance the compressive mechanical stress generated by MHS. Importantly, the model reveals that epithelia cannot be treated as ideal semi-permeable membranes. Instead, cross-coupling between ion and water fluxes that arise from paracellular pathways (66) and co-transport (67) processes must be considered. While a cell-level description of hydraulic processes is needed to capture cell shape changes during MHSs compression, the present framework already establishes a quantitative basis for connecting cell-level transport activity to emergent tissue-scale mechanics and offers a platform for predicting how epithelial geometry and density influence mechanical stress generation.

In vivo, epithelia in organs such as the kidney and intestine routinely handle large trans-epithelial fluxes (68, 69). Our findings imply that these transport processes may also contribute to the mechanical regulation of tissue homeostasis, complementing or even surpassing the role of contractility in certain contexts. We thus identify active ion transport as a powerful source of epithelial mechanical stress that enables tissues to achieve and maintain density homeostasis. By shifting the focus from contractility to hydraulics, our work expands the conceptual framework of epithelial mechanics and points toward fluid transport as a key regulator of tissue behavior.

## Materials and Methods

### Cell lines origin and maintenance

The MDCK cell lines MDCK Histone H1 GFP (70), MDCK LifeAct GFP (60), MDCK NMMIIA KD (44) and MDCK ZO-1/2 dKD (51) were used. To obtain MDCK CAAX-GFP cells, MDCK cells were transfected by electroporation using NeonTM Transfection System (Thermo Fisher, 100 μL reaction kit, Ref MPK10096, 1650 V, 20 ms, 1 pulse). The stable MDCK CAAX-GFP clone was selected after two rounds of fluorescence-activated cell sorting (FACS). All MDCK cell lines were cultured in Dulbecco’s modified eagle medium with 4.5 g/L glucose and L-glutamine (DMEM, Gibco, 31966-021) and 1% penicillin-streptomycin (Gibco, 14140-112), containing 10% fetal bovine serum (BioWest, Cat# S1810-500), (denoted as DMEM-FBS) media at 37 °C incubators supplemented with 5% CO_2_. Caco2 cells (from ATCC) was cultured at 37 °C incubators supplemented with 5% CO_2_ in the same DMEM medium but containing 20% FBS.

### Fabrication of micro-sized hydrogel spheres (MHSs)

The MHSs were fabricated using a water-in-oil emulsion protocol as summarized in Figure S1a, 20 µL aqueous PEGDA (mw. 700 Da, Sigma, Cat.455008) solution containing 10% PEGDA (MHS), 7% (for MHS_soft) or 15% (for MHS_stiff), 0.3 mg/mL photo-initiator Irgacure^@^ 2959 (BASF. Cat.55047962), 0.2% surfactant Sodium dodecyl sulfate (SDS, Euromedex, EU0660), 0.05 mg/mL fibronectin (Merck Millipore, FC010) and 0.2 mM acryloyloxyethyl thiocarbamoyl rhodamine B (Acryl-RhoB, Polysciences, Cat.25404) was added to 1 mL mineral oil (Sigma, Cat.M8410), followed by 1 min of vortex and 30 s of UV illumination. Then, 2 mL DMEM-FBS medium was added to the emulsion with subsequent centrifugation at 1000 rpm for 3 min to precipitate the formed MHSs. Several DMEM wash and centrifugation processes were conducted to completely remove the mineral oil, then the MHSs were kept in DMEM medium and incubated at 37 °C until being used.

### Cell seeding on MHSs

After trypsin (Gibco, 25300-054) treatment, cells are centrifuged and re-suspended in DMEM-FBS, then the cells are mixed with prepared MHSs at a desired cell density in a low adhesion plate. After 1-2 hours of incubation, the MHSs are washed with fresh DMEM_C medium to remove the non-adhered cells.

### PAA gel fabrication

21 kPa Polyacrylamide gels (PAAm) were prepared as described previously (73). The PAAm gels were washed with 10 mM HEPES and coated with a 50 *μ*g/mL fibronectin solution prior to cell seeding.

### Drug treatments

During time-lapse imaging process, warm DMEM-FBS containing certain drug (first dissolved in DMSO at high concentration, and then diluted 1000 times with culture medium to the final concentrations as indicated in Table S1.) was added into the live imaging sample after removing the pure DMEM-FBS at certain time point. All those manipulations were conducted in between the acquisition intervals, therefore imposed no change to the registered data points and acquisition settings. The details of used drugs can be found in Table S1.

### Live imaging

Cell-laden MHSs are resuspended in DMEM-FBS, then mixed with 1% liquefied agarose gel at a temperature around 37 °C quickly and gently with a volume ratio of 1:1, then poured to a glass-bottom petri dish and allowed the gelling of agarose gel at room temperature (RT) for 3-5 min before adding DMEM-FBS. The prepared samples are incubated at 37 °C for at least 2 h before live microscopic observations. Time-lapse live images were performed either on a multi-channel inverted microscope (Olympus, IX83) equipped with temperature and CO_2_ control box or on a high-resolution spinning disk microscope (Nikon, Ti2 Eclipse) equipped with temperature and CO_2_ control box. Multiple (x, y) positions, z stacks (1-3 µm per step, 12-120 steps in total) and multi-channels were utilized to obtain the precise MHS sizes and/or cellular activities.

### MHSs deswelling experiments

Fabricated MHSs were first encapsulated in 0.5% agarose gel and incubated in DMEM-FBS at 37 °C for more than 24 hours to reach an equilibrium swelling state. Then the MSHs were imaged with high resolution spinning disk microscope (Nikon, Ti2 Eclipse) under the condition of live cell imaging to reveal the original MHS sizes. Then, the sample is rinsed more than 3 times using DMEM-FBS containing extra NaCl of various concentrations, followed by a more than 24 hours incubation to ensure equilibrium swelling of the MHS at 37 °C before spinning disk imaging. The incubation of MSHs in different media followed a fashion where NaCl concentration changed from low to high (1-5+ M), and the obtained sizes were normalized by the original MHS size.

### Osmotic shock experiments

MDCK cells were seeded on MHSs then under live-imaging in normal osmolarity conditions. After the MHSs were deformed, pre-warmed new medium with varied osmolarities was added into the samples after carefully removing the old medium. The shocks were applied in between acquisition intervals therefore no other experimental parameters were modified except for medium osmolarity. To prepare osmotic shock media, 100 mM sucrose and 150 mM NaCl was added to physiological culture medium (300 mM) to obtain hypertonic shock media of 400 mM and 450 mM, respectively; 1 mL, 1.5 mL and 2 mL Milli Q water was added to 2 mL, 1.5 mL and 1 mL physiological culture medium to obtain 200 mM, 150 mM and 100 mM hypotonic shock media, respectively.

### Indentation experiments

2D PEGDA gels with the same formulation as PEGDA MHSs were equilibrized in DMEM_C media before indentation were performed with a nanoindentor (Chiao, Optics11 Life) mounted on an inverted multi-channel microscope (Olympus, IX83) at room temperature, with a spherical tip of 9 µm and a cantilever stiffness of 0.53 N/m. The indentation depth was set at 1.5 µm and acquired data were fitted with Hertz model to obtain the Young’s modulus E. At least 3 parallel samples were measured for each rigidity. At least 10 static measurements on different regions of the sample were performed for each sample.

### Compressive stress evaluation

The compressive stress (Δ*P*) required to deform isotropically a homogenous material can be evaluated from the bulk modulus (*B*) of the material using the following relation (71):

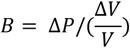

where *V* is the initial volume of the material and Δ*V* is the volume change of the material. In our experiments, the bulk moduli B of the MHS exhibiting two different rigidities are calculated according to the following relation:

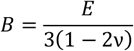

where *E* is the Young’s modulus (obtained from indentation experiments), and *ν* is the Poisson ratio. From a comprehensive study by J. Cappello et al. (72) we estimate here for the MHSs of both rigidities exhibit a *ν* ≃ 0.25.

### Volumetric flux calculation

The average volumetric flux or water flux rate per unit area *q* was estimated from the MHS compression as follows:

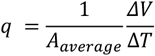

where *ΔV* is the volume loss of the MHS during the duration Δ*T* of the deformation, and *A*_*average*_ is the averaged compression-onset and compression-end surface area of MHS.

### Immunostaining and confocal imaging

Cell adhered MHSs at different time points/ tissue growing stages are fixed and stained by immuno-fluorescence with a modified paraformaldehyde (PFA) protocol, details of used agents and antibodies can be found in Table S1.

Briefly, samples embedded in agarose gels were fixed with 4% PFA in PBS (30 min, room temperature), washed 3 time for 3-5 min with PBS then stored in PBS at 4 degrees overnight. Next day, the samples were taken out and let to warm up to room temperature. Then the samples were permeabilized with 0.5% Triton in PBS (30 min, room temperature) and blocked (1% BSA in PBS) for 2 hours followed by 15 min wash in PBS. Samples were subsequently incubated overnight at 4 °C with primary antibodies in blocking buffer under agitation. Next day, they were washed with PBS for 3 times, 10 min each. Samples were then incubated with secondary antibody at room temperature for 3 hours under agitation, followed by extensive washing in PBS. Then the samples were mounted with Vectashield mounting media on cover slides and sealed with dental glue for imaging, using either a Zeiss LSM 980 Airyscan confocal microscope with glycerin immersion objectives (25x or 40x) at 0.5 to 1 *μ*m per stack, or a Nikon Ti2 Eclipse spinning disk confocal microscope with 20x air objective at 0.3 to 1 *μ*m per stack.

### Data analysis

All the images (including time-lapse live image) obtained were either analyzed by Fiji, Imaris (10.1.1) or CellPose2. The statistics obtained from image analysis and other measurements were processed either by Origin (2017) or GraphPad (Prism 9 or 10). Sigmoid fitting analysis was performed first using MATLAB and further processed by GraphPad (Prism 10). Unless otherwise indicated, all plots show the Mean ± SD. P-values were calculated by t test, ns p > 0.05, * p < 0.05, ** p < 0.01, *** p < 0.001, **** p < 0.0001.

Specifically, segmentation of confocal images showing cell apical membrane marker (either MDCK CAAx-GFP or E-cadherin GFP) were conducted using CellPose2, then cell areas were quantified according to the segmentation using Fiji. Note that the curvature of MHS wasn’t taken into account. Cell heights were quantified manually using Fiji tools, the highest point of a cell was extracted over a whole z-stack image. Cell displacement on the MHSs was manually quantified over live cell imaging videos, note that the curvature of the MHS wasn’t taken into account, thereby expressing underestimated cell velocities.

## Author Contributions

RMM and BL conceived and supervised the project. HW, TD, GA, RC carried out the experimental investigations. HW, CD, JP, BL, RMM analyzed the experimental results. CD and JP developed the theoretical model. All authors contributed to discussions. HW, CD, JP, BL and RMM wrote the manuscript.

## Acknowledgements

We would like to thank Joseph d’Alessandro for some codes, as well as all the members of the “Cell adhesion and Mechanics” team for helpful discussions. This work was supported by the European Research Council (Grant No. Adv-101019835 to BL), LABEX Who Am I? (ANR-11-LABX-0071 to BL and RMM), the Ligue Contre le Cancer (Equipe labellisée 2019 to RMM), the Alexander von Humboldt Foundation (Alexander von Humboldt Professorship to BL), Institut National du Cancer (INCa_16712 and INCa_18429 to BL and RMM), the Agence Nationale de la Recherche (“STRATEPI” DFG-ANR-22-CE92-0048) to RMM). We acknowledge the ImagoSeine core facility of the IJM, a member of IBiSA and France-BioImaging (ANR-10-INBS-04) infrastructures.

## Data availability

All data is included in the manuscript and/or supporting information

## Supplementary figures and Tables

**Figure S1.**
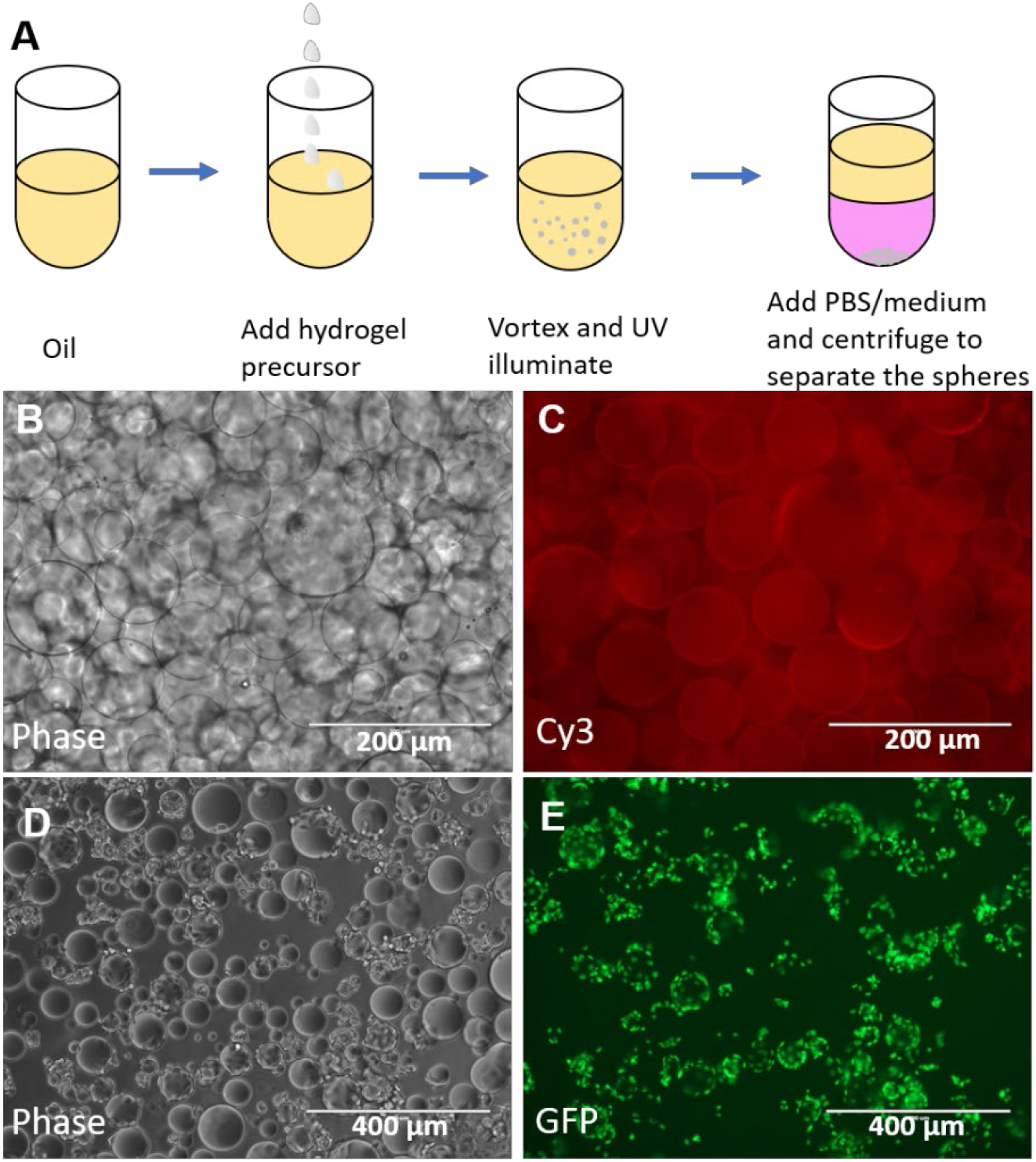
MHS fabrication and epithelial cell seeding. **(A)** MHSs fabrication procedure. **(B)** Phase contrast image of fabricated MHSs and **(C)** corresponding cy3-laballed fibronectin signal. **(D)** Phase contrast image of MDCK-Histone GFP laden MHSs and **(E)** corresponding nuclear signal image.

**Figure S2.**
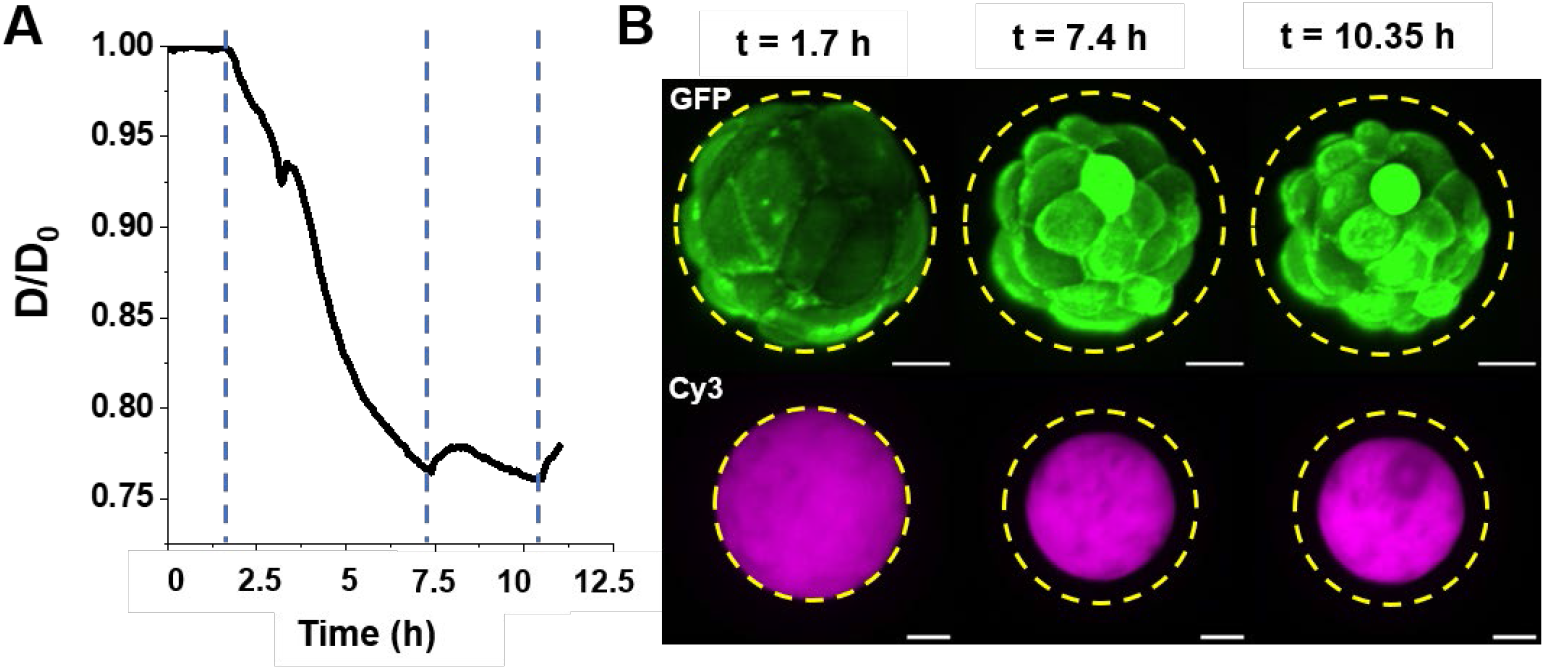
Water loss in the MHS-epithelium system. (**A**) Representative evolution curve of MDCK LifeAct GFP cells on MHSs. **(B)** Corresponding full z projection of GFP channel (cells) and max-z projection of Cy3 channel (MHS) images at different time points indicated by the blue dot line in (A), the yellow dot circles mark the initial outer diameter of the MHS-cells system (GFP channel) and the MHS size (Cy3 channel) at the onset of MHS deformation. Scale bars = 20 µm.

**Figure S3.**
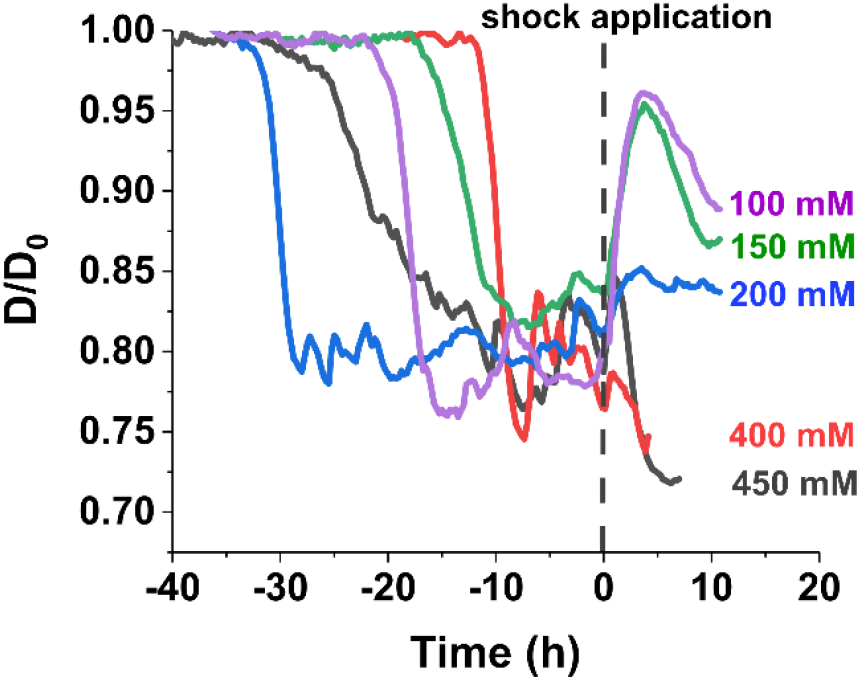
Osmotic shock. Representative size evolution curves of cell covered MHSs under osmotic shocks, here the curves were normalized to the shock application time t=0, mM = mmol/L.

**Figure S4.**
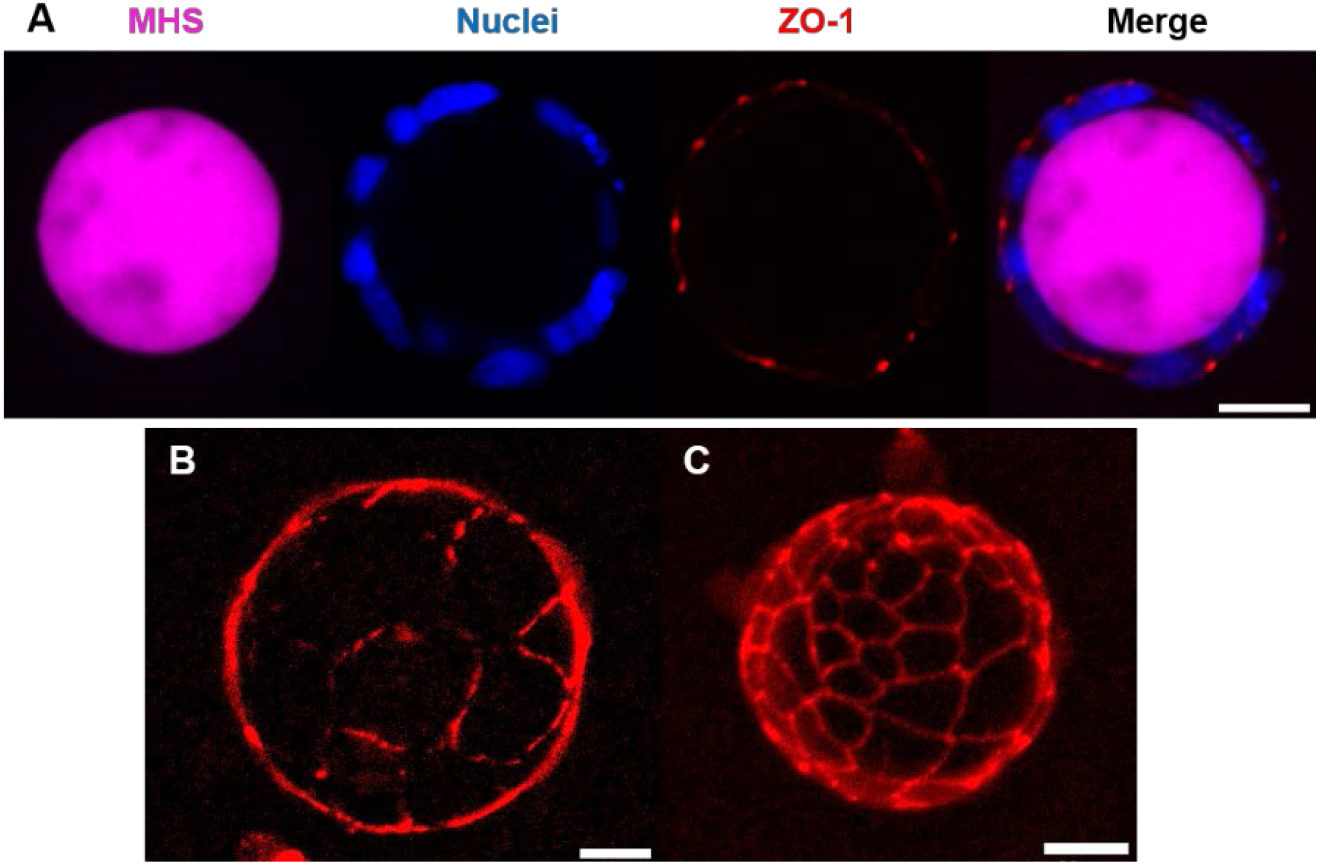
Immunostaining of ZO-1. (**A**) ZO1 labelled tight junction are located at apical position (facing toward the culture medium) in between each cell. Hemisphere z-stack projection of ZO1 staining (**B**) before MHS and (**C**) after MHS deformation. Scale bar = 20 µm.

**Figure S5.**
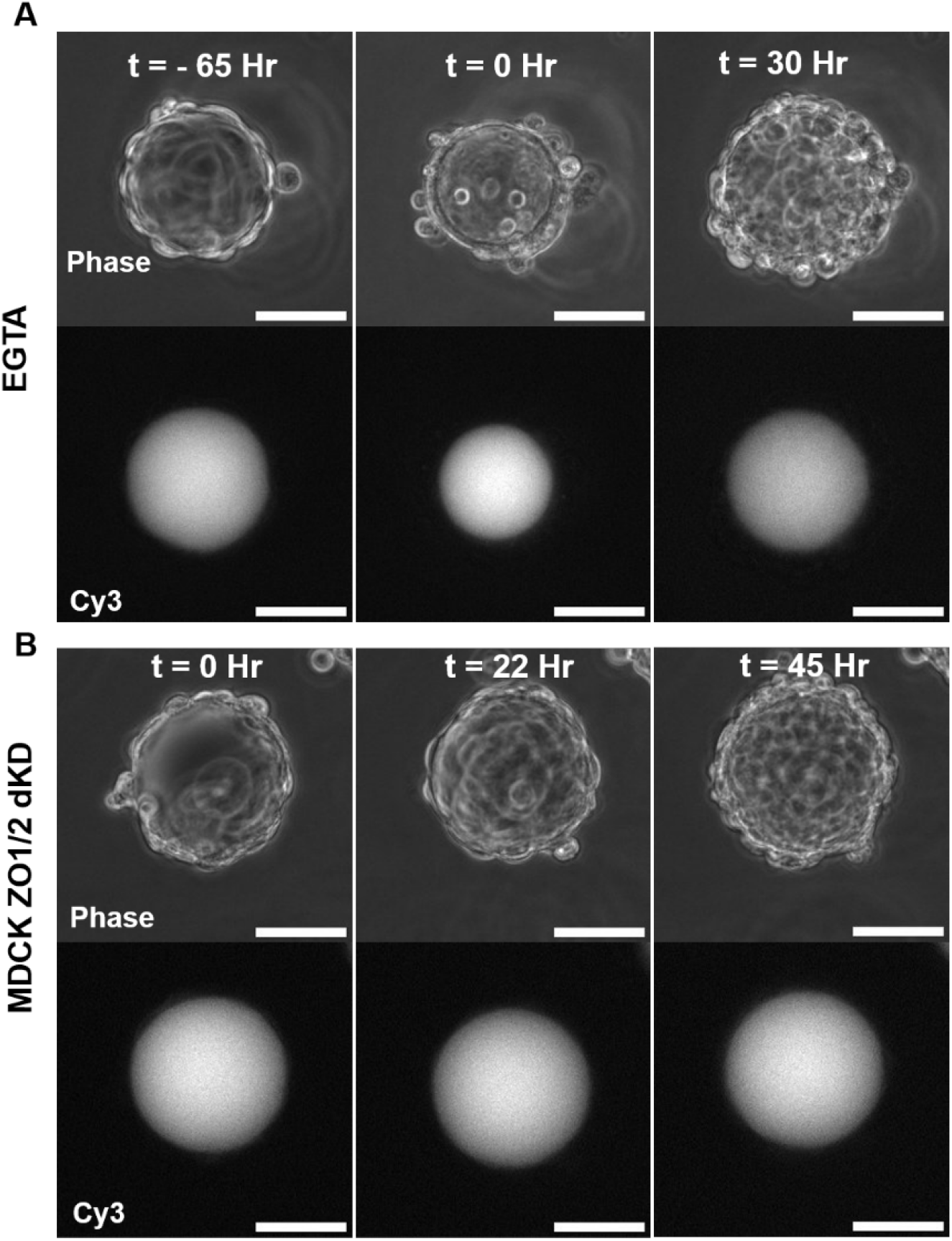
Images of epithelial tissue barrier function modulation. **(A)** Phase images and cy3 channel of MHS at t = -65 hr, t= 0 hr and t = 30 hr, which correspond to the curve Fig.2B. **(B)** Phase images and cy3 channel images of MHS at t = 0 hr, t = 22 hr and t= 45 hr, which correspond to a representative curve of MDCK ZO-1/2 dKD cell seeded MHS. Scale bars = 50 µm.

**Figure S6.**
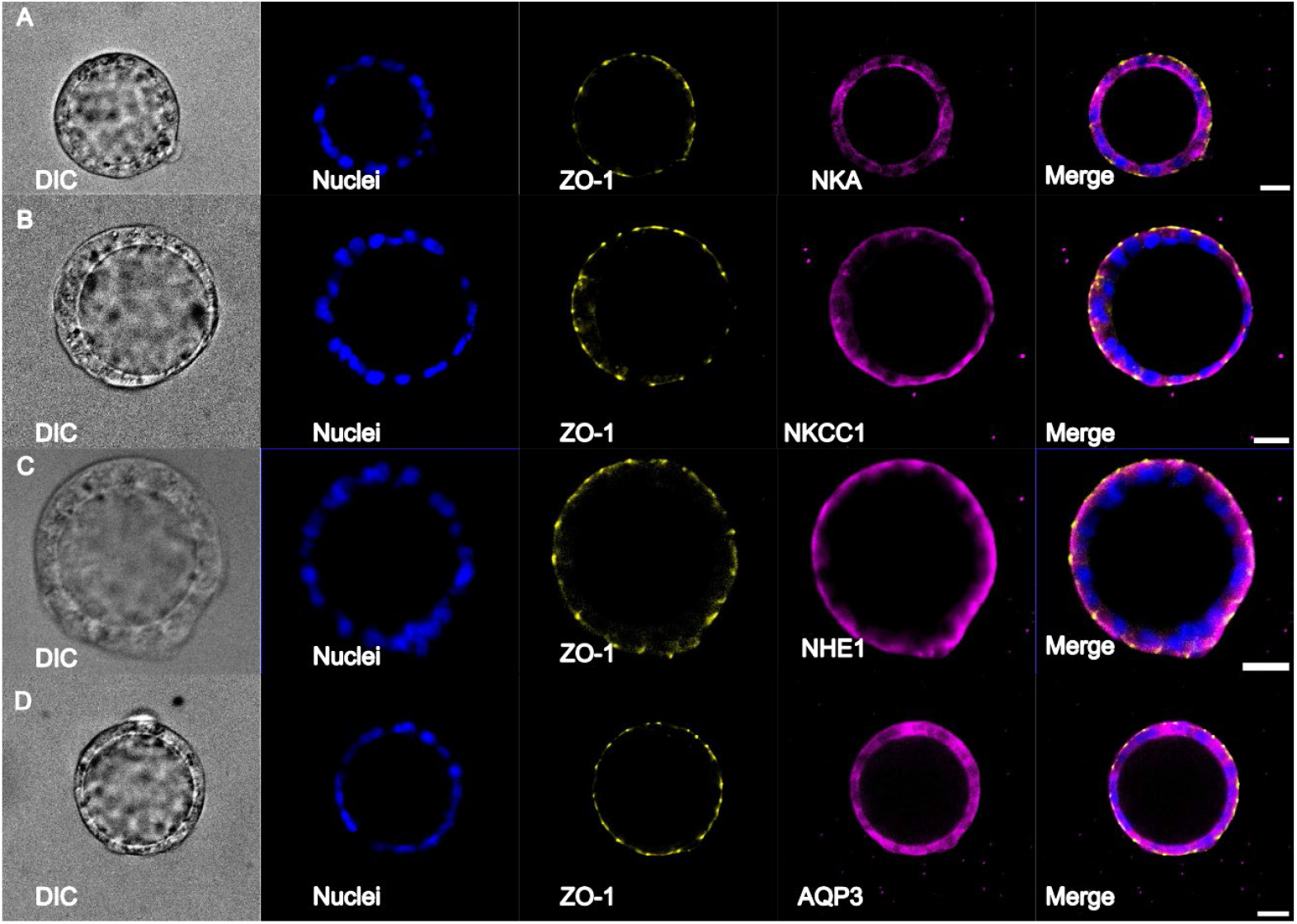
Immunostaining of ion pump/channels and water channels, together with ZO-1 staining marking cell apical domain. **(A)** Immunostaining of NKA pump shows a basal-lateral distribution. Immunostainings of NKCC1 **(B)** and NHE1 **(C)** show an apical distribution. (**D**) AQP3 immunostaining shows no preferential distribution. Scale bars = 20 µm.

**Figure S7.**
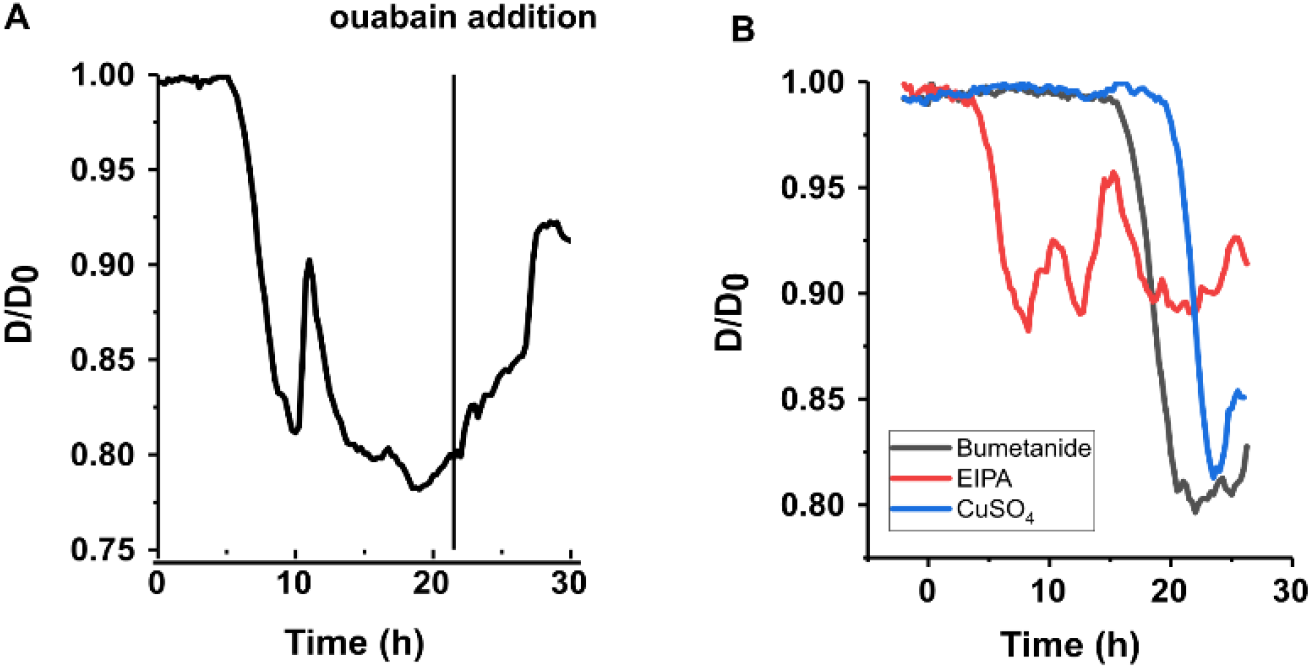
Representative MHS size evolution curves under ion pump/channel inhibition conditions. **(A)** Ouabain addition results in partial MHS size recovery. **(B)** Representative MHS size evolution curve under inhibitions of bumetanide, EIPA and CuSO4, respectively. Drugs were added at t = 0h for all groups.

**Figure S8.**
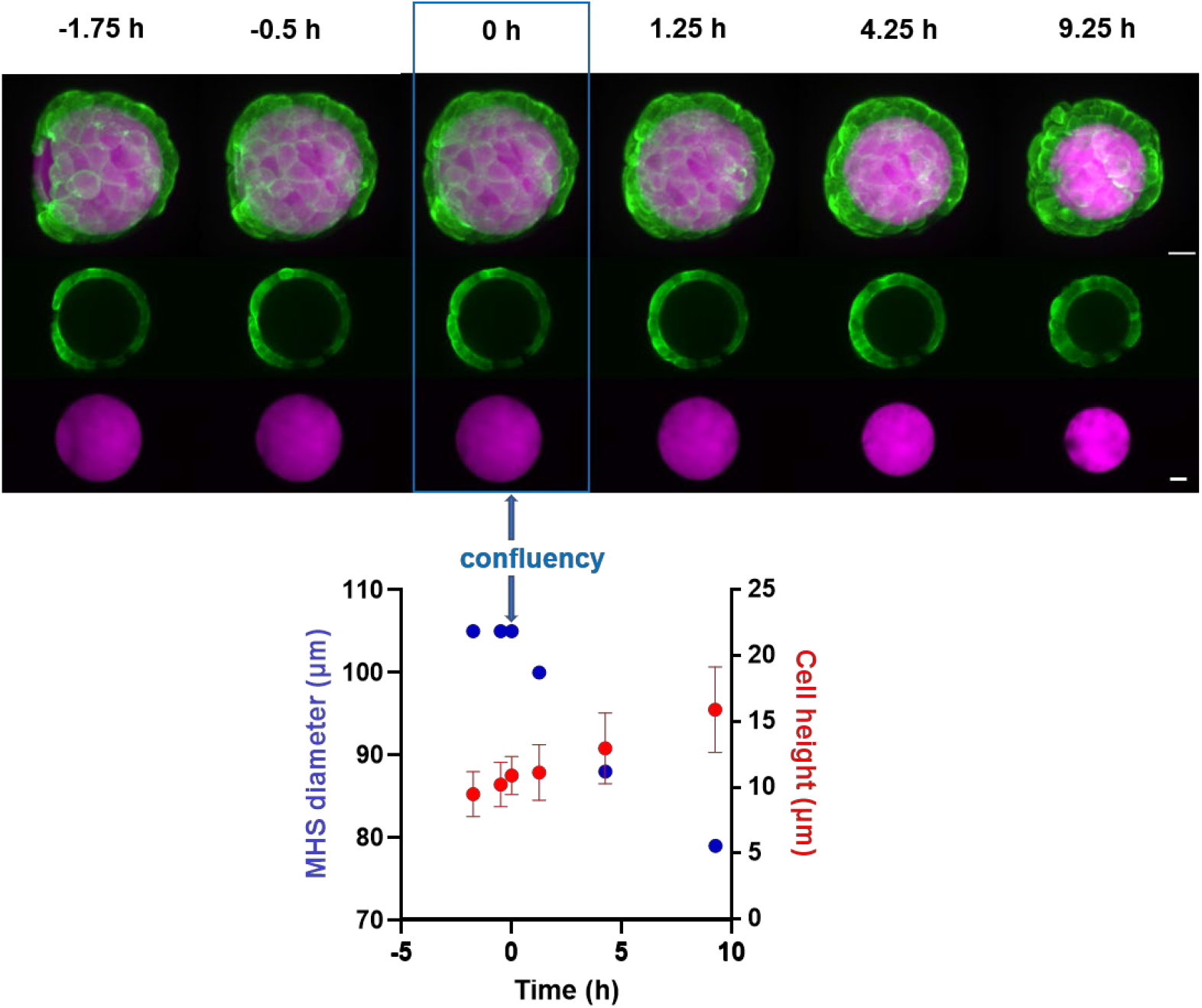
Cell height evolution and corresponding images. Representative cell height evolution quantification and corresponding image during a live lapse course. Here t= 0h is the confluency. Scale bars =20 µm.

**Figure S9.**
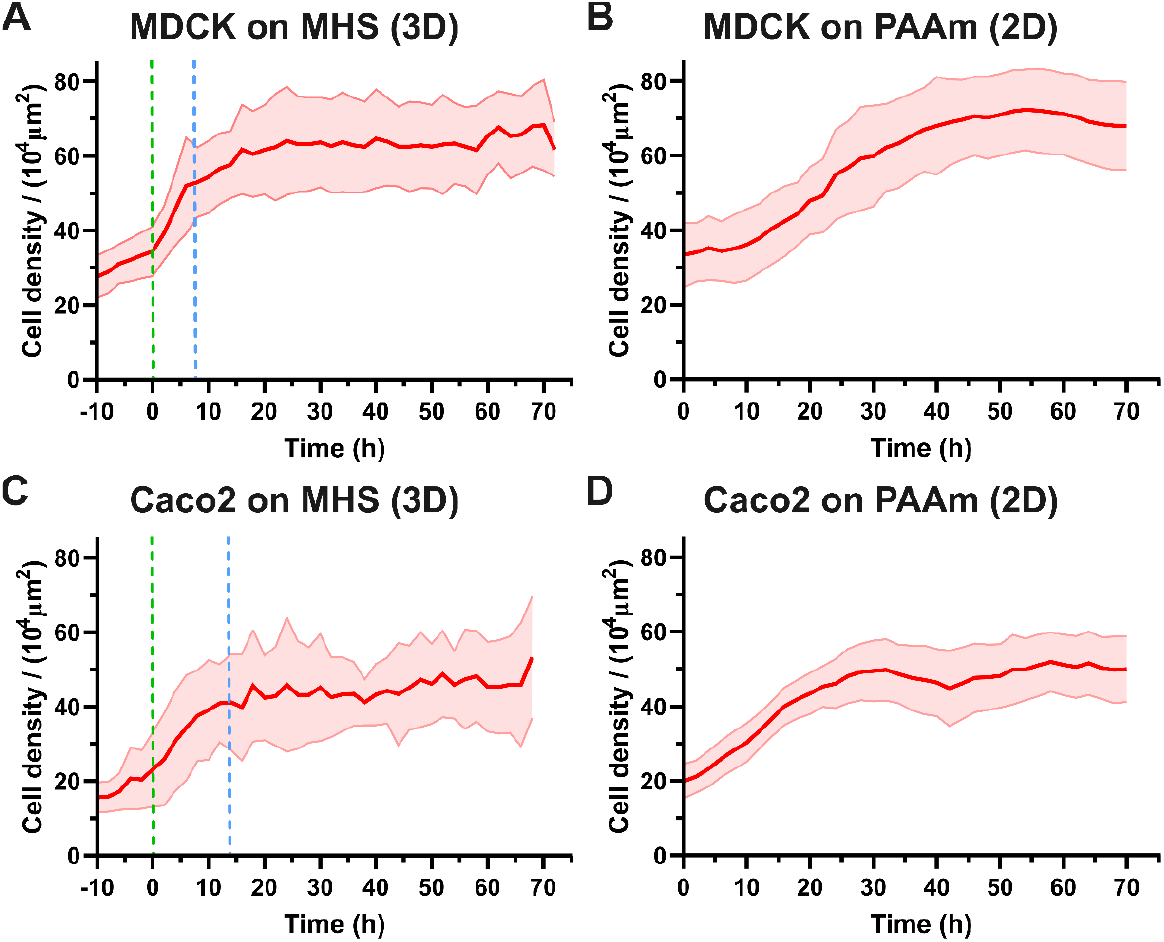
Cell density evolution of MDCK and Caco2 on MHSs and 2D PAAm. (**A**) Cell density of MDCK on MHSs (n=8, N=1) (**B**) and on 2D PAAm 21kPa substrate over time (n=3, N=1). (**C**) Cell density of Caco2 on MHSs (n=6, N=1) (**D**) and on 2D PAAm 21 kPa substrate over time (n=2, N=1). For cells on MHSs, the dotted green and blue lines represent respectively the onset and end of MHSs compression. The origin of time for cells on 2D PAAm substrate corresponds to the onset of confluence in the field of view.

**Figure S10.**
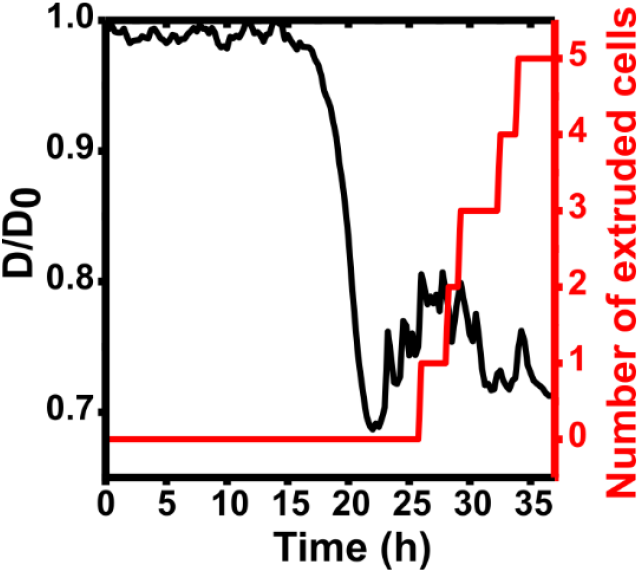
Representative curves of MHS size evolution and sum of the extruded cell number as a function of observation time.

**Figure S11.**
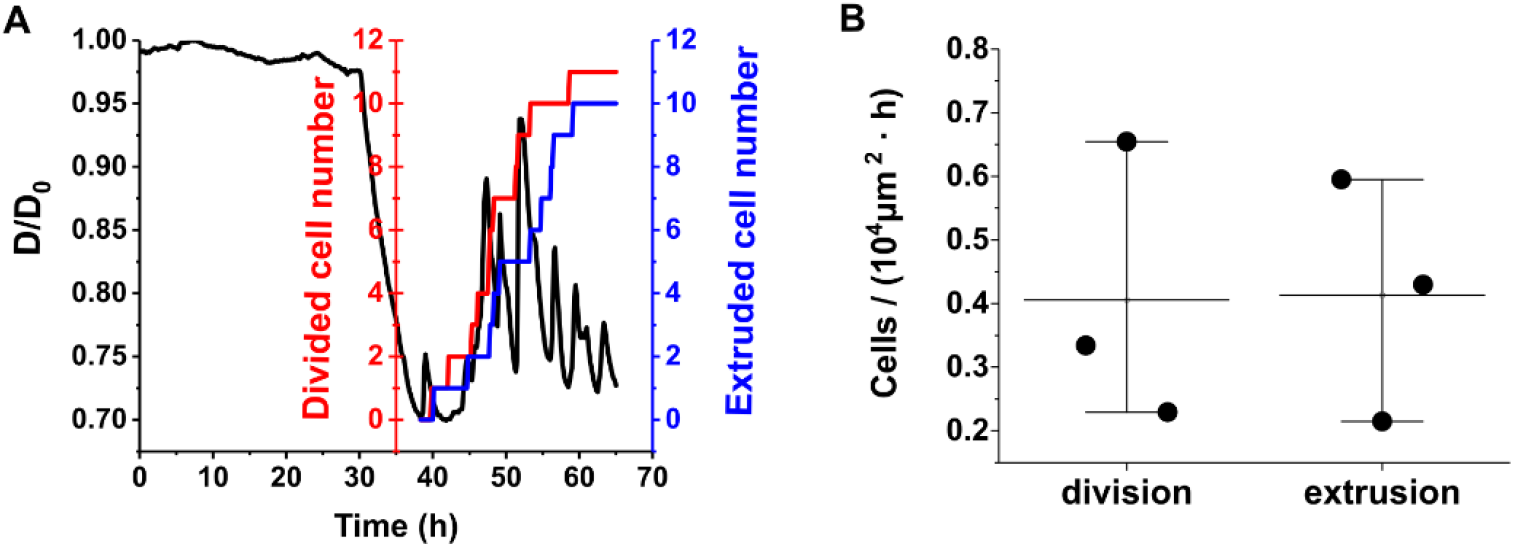
Balance of cell extrusion and division post MHS compression. (**A**) Representative curve of cumulative cell division and cell extrusion event with time after MHS compression (quantification issued from Video S4). (**B**) Quantification of cell division and cell extrusion frequency after MHS deformation, n = 3, N = 2.

**Figure S12.**
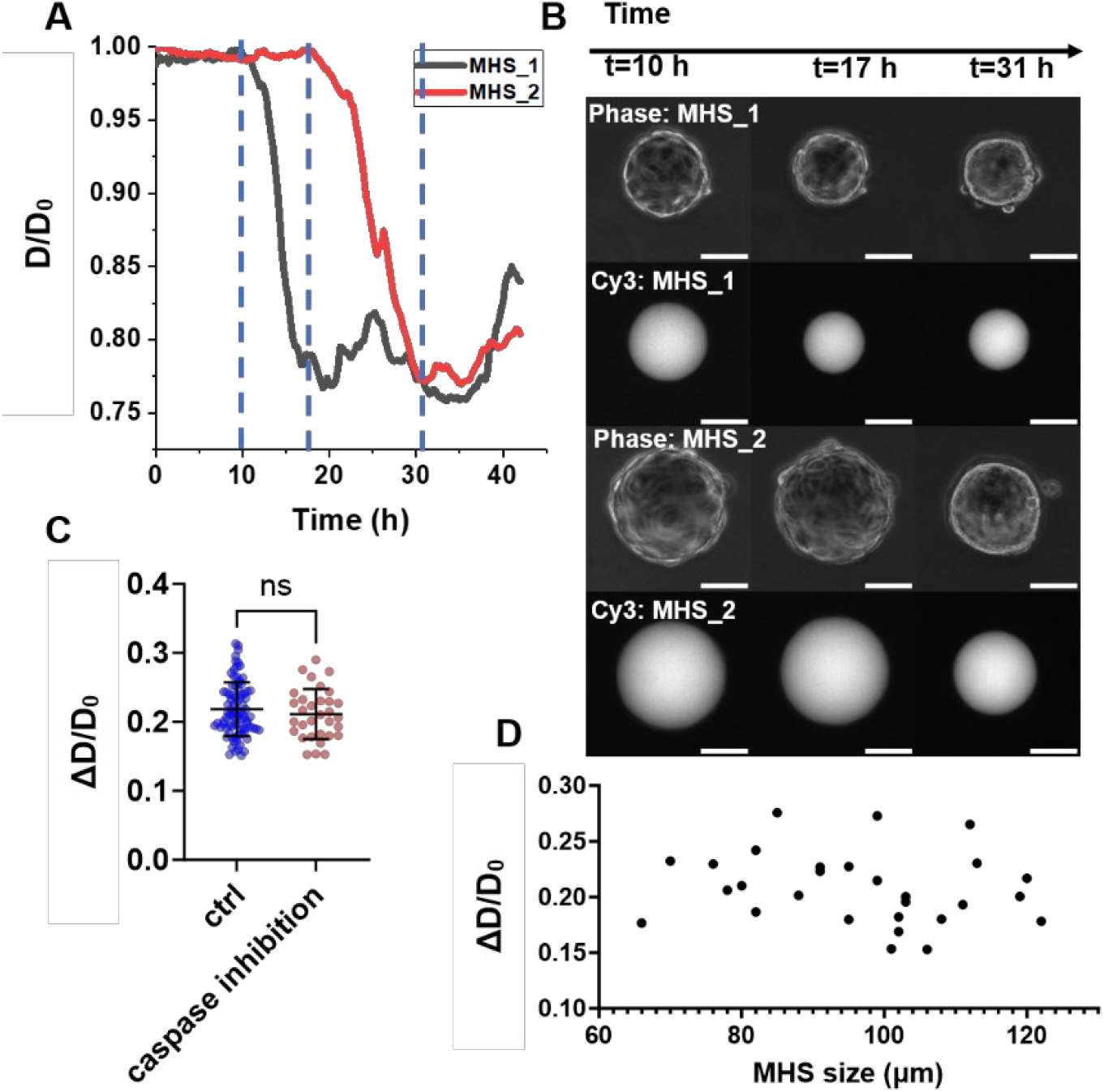
MHS deformations under apoptosis inhibition conditions. **(A)** Representative evolution curves of two MHSs, MHS_1 and MHS_2 under apoptosis blockage. **(B)** Corresponding phase contrast (cells on the MHS) and Cy3 channel (MHS) images at different time points indicated by the blue dot line in (A), scale bars = 50 µm. **(C)** Quantification of maximum deformations of MHSs under apoptosis inhibition does no show significant difference when compared with control group. **(D)** The maximum deformations of MHS as a function of MHS size, showing no obvious size dependency when under apoptosis inhibition.

**Figure S13.**
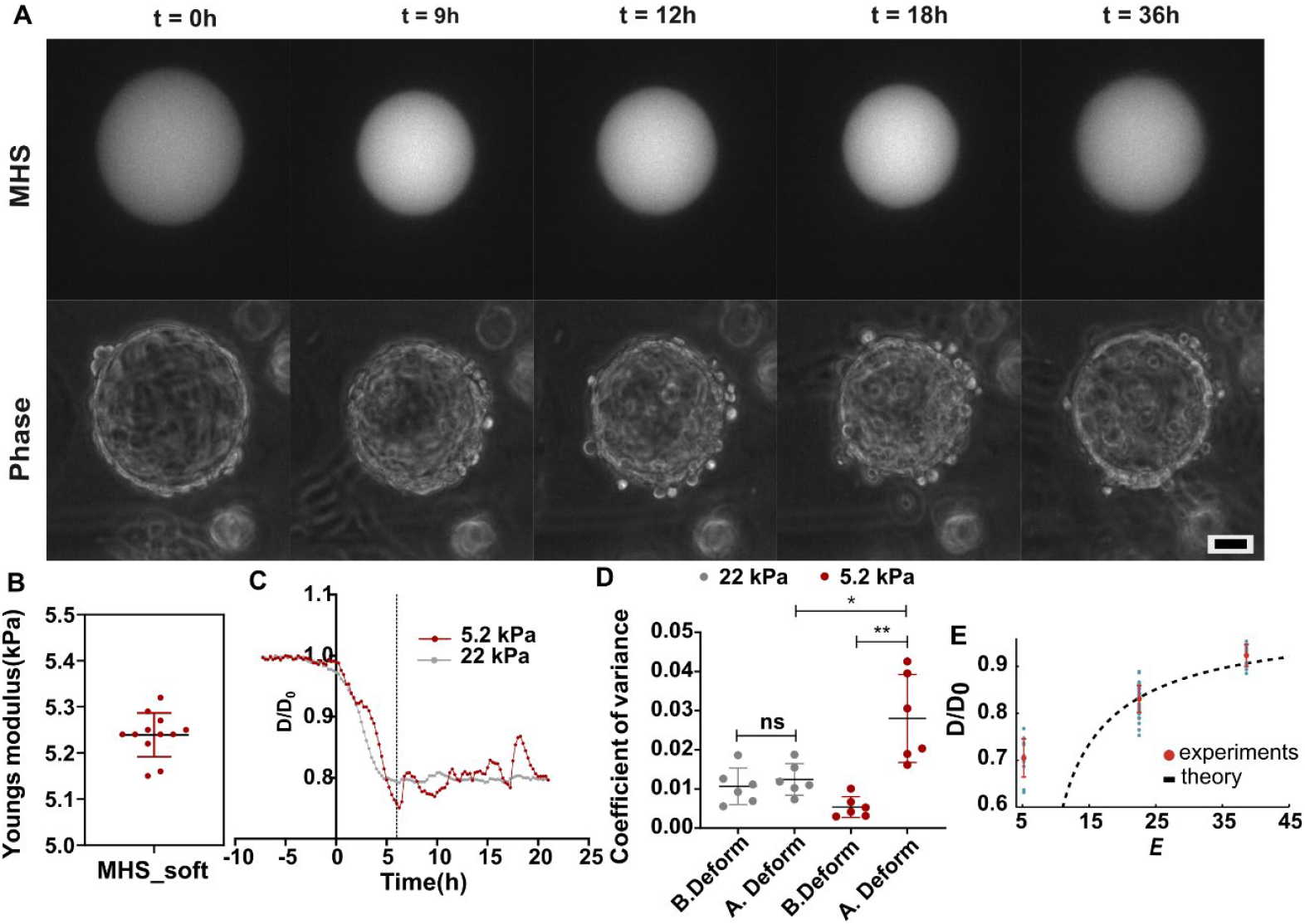
MDCK on 5.2 kPa MHS. **(A)** Representative time-lapse images of ∼5.2 kPa MHS, showing the fluctuations of MHS size, with Phase image, 0 h marking the onset of compression (scale bar, 25 μm). (**B**) Stiffness of the bead for 5% PEGDA (**C**) Normalized bead diameter (D/D0) plotted over time shows pronounced post-compression fluctuations in ∼5.2 kPa MHS relative to ∼22KPa MHS. (**D**) Coefficient of variance (*CV* = σ/ μ, σ is sample standard deviation and μ is sample mean) of D/D0 reveals that soft MHS (∼5.2 kPa) fluctuate (0.0280 ± 0.0046) significantly more after compression, whereas stiffer MHS (∼22 kPa) remain stable (0.0124 ± 0.0017) (*two-tailed Student’s t test; *P < 0.05, **P < 0.001, n = 6 MHS per condition). (**E**) Normalized steady-state radius D/D_0 as function of the gel Young’s modulus E (in kPa). The black dashed line is a prediction from Eq. (4) of the main text and using the parameters fitted in Fig. 5C.

**Table S1.**
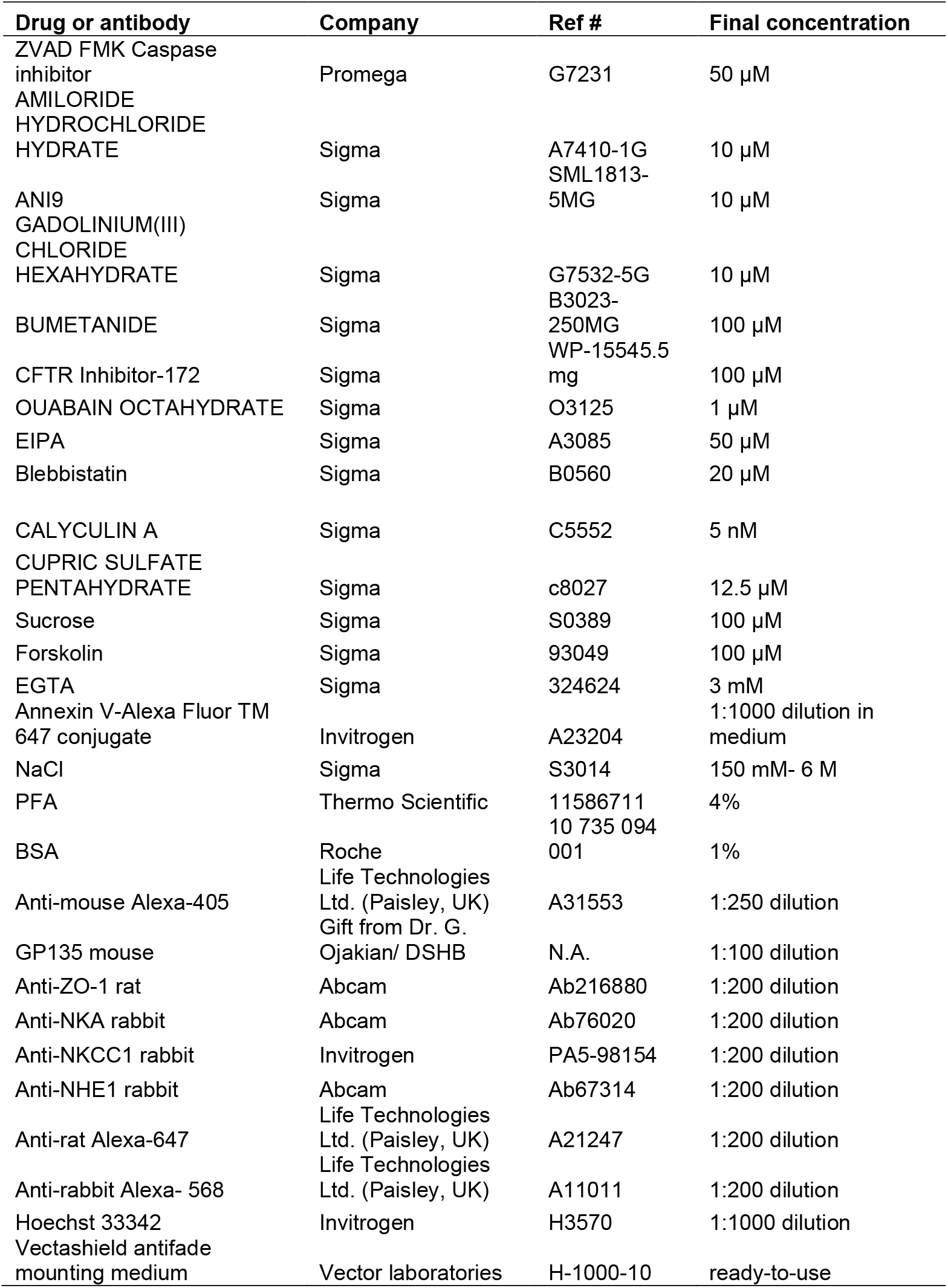
Agents and antibodies list for the experiments

## Supplementary information

### I. DETERMINISTIC MODEL

#### A. Force balance and water transport

We consider a simple model for a cell spheroid compressing a soft gel. The cell spheroid is described as a permeable sphere of radius *R*(*t*) enclosing the soft gel with compression modulus *B*.

##### Force balance

Calling *P* ^in,out^ the pressure inside and outside the spheroid, the mechanical balance reads:

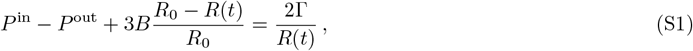

with Γ the spheroid surface tension and *R*_0_ = *R*(*t* = 0) the spheroid radius before compression.

##### Volume conservation

Fluid exchanges also dictate the spheroid volume. The lumen volume *V* = (4*/*3)*πR*(*t*)^3^ is controlled by water influx and its dynamics reads [1, 2]:

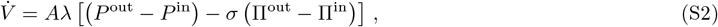

where 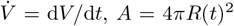 denotes the spheroid surface area, *λ* is the membrane permeability to water flows, and Π^out^ − Π^in^ ≃ − *k*_B_*T* Δ*C* with Δ*C* = *C*^in^ − *C*^out^ denotes the osmotic pressure difference between the outside and inside. Importantly, we have included a so-called reflection or selectivity coefficient *σ* [1, 3]. A fully semi-permeable membrane corresponds to the case *σ* = 1, while a membrane that would be fully permeable to water and to osmolytes would have *σ* = 0.

##### Ion conservation

For simplicity, we consider a single ionic species. The number *N* ^in^ of ions in the lumen reads [2]:

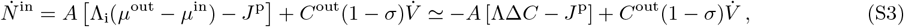

where *µ*^in,out^ = *k*_B_*T* log *C*^in,out^ are the chemical potentials inside and outside the spheroid. We have also introduced the Onsager coefficient Λ_i_ and a contribution *J*^p^ to the flux due to active pumps (positive when the flux is outwards). To obtain the second equality of Eq. (S3), we have considered a small concentration difference Δ*C* to expand the log and we have introduced the ion permeability Λ = Λ_i_*k*_B_*T/C*^out^. Note finally that since Δ*C* = *N* ^in^*/V* − *C*^out^, and considering that the concentration outside the spheroid is constant, we deduce:

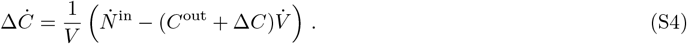

Equations (S1)-(S4) can then be rewritten as two coupled equations:

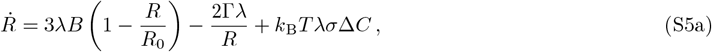

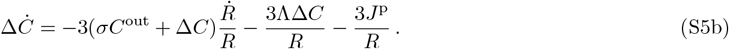

from which we can deduce the steady-state normalized radius:

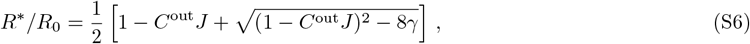

where *J* = *σJ*^p^*/*(3*B*Λ_i_) and *γ* = Γ*/*(3*BR*_0_).

#### B. Dimensionless equations and state diagram

##### Dimensionless equations

We introduce the dimensionless variables:

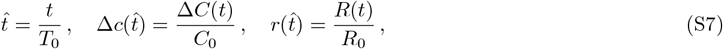

*t*such that Eq. (S5) become:

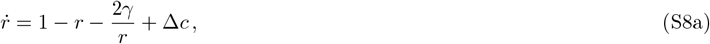

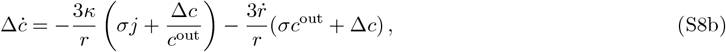

where we introduced the reference parameters

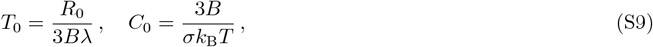

and the dimensionless parameters:

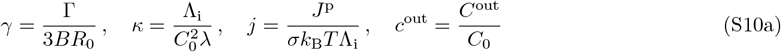

##### Steady-state solution and linear stability analysis

The steady-state radius and concentration difference are:

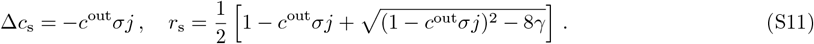

The stability of these steady-state solutions can be obtained by looking at the dynamics of a perturbation *r*(*t*) = *r*_s_ + *δr*(*t*), Δ*c*(*t*) = Δ*c*_s_ + *δc*(*t*). We find:

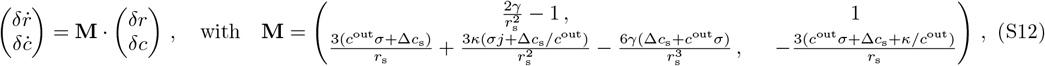

and the stability is given by the real part of the eigenvalues of the stability matrix **M**. Equation (S11) and the eigenvalues obtained from the stability matrix were used to obtain the state diagram given in the main text. The parameters used are: *κ* = 0.6, *σ* = 0.04, *c*^out^ = 1.5.

#### C. Orders of magnitude

We now estimate the values of the dimensionless parameters of the model. The experiments provide an initial sphere radius *R*_0_ ≃ 50 *µ*m, a compression modulus *B* ≃ 20 kPa (soft gel) and an initial concentration *C*^out^ ≃ 300 mmol/L. From the literature, we can estimate the tissue surface tension Γ ≃ 10^*−*3^ − 10^*−*2^ N/m [4], hydraulic permeability *λ* ≃ 10^*−*14^ − 10^*−*13^ m/(Pa·s) [5], ion permeability Λ = Λ_i_*k*_B_*T/C*^out^ ≃ 10^*−*9^ − 10^*−*6^ m/s [2, 6], and active ion flux *J*^p^ ≃ 10^17^ − 10^18^ m^*−*2^·s^*−*1^ [2]. We thus obtain order-of-magnitude estimates for the dimensionless parameters:

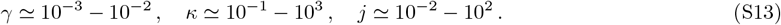

The parameters used for the simulations below are consistent with these estimates.

#### D. Role of diffusion

In the model presented above, the concentrations inside the gel and in the surrounding medium are both considered homogeneous, and the effect of diffusion was neglected. We discuss here the limits of this assumption.

Considering a diffusion constant for ions to be *D* ≃ 2000 *µ*m^2^*/*s and a sphere radius of *R*_0_ ≃ 50 *µ*m, the typical time *τ*_in_ to reach a homogeneous concentration within the gel is of order 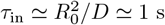 s. It is therefore reasonable to consider that concentration inside the gel is homogeneous. Outside the sphere, the total volume of the medium is *V*_tot_ ≃ 3 mL, from which we deduce the typical time *τ*_out_ to reach a homogeneous concentration outside to be of order *τ*_out_ ≃ *V* ^2*/*3^*/D* ≃ 42 h. This indicates that the hypothesis of a homogeneous concentration outside the sphere is not reasonable. However, we show below that taking into account diffusion outside the sphere leads to slightly faster compression but does not qualitatively change the conclusions of the model.

##### Diffusion in flat geometry

To discuss the role of diffusion we consider a flat 1d geometry for simplicity. The total length of the system is *L*, and the gel is located between *x* = 0 and *x* = *h*(*t*), while the space between *x* = *h*(*t*) and *x* = *L* is the external medium. The epithelium is located at the interface between the gel and the outer medium.

##### Solution of the diffusion equation

We first solve the diffusion problem in the outer region *h* ≤ *x* ≤ *L*. Up to a translation and considering *L* ≫ *h*, we thus have to solve the following problem:

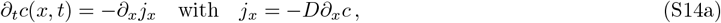

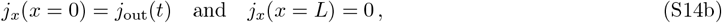

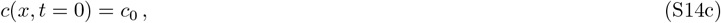

with *c*(*x, t*) the concentration in the outer medium, *D* the diffusion constant and *j*_out_(*t*) the time-dependent ion flux from the epithelium.

A solution of Eq. (S14) can be written as:

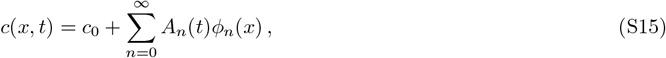

where the *ϕ*_*n*_ are the eigenfunctions of the Laplacian and satisfy 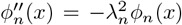 with 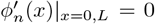, which yields *ϕ*_*n*_(*x*) = cos *λ*_*n*_*x* and *λ*_*n*_ = *nπ/L*. These eigenfunctions are orthogonal:

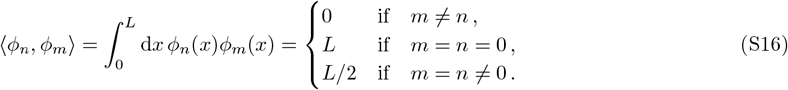

Differential equations for the coefficients *A*_*n*_(*t*) can then be obtained by computing 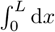 (S14a) ×*ϕ*_*m*_(*x*) and using the scalar product defined above. We obtain:

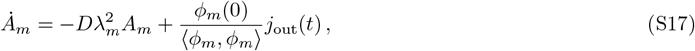

with *ϕ*_*m*_(0) = 1.

##### Hydraulics with diffusion in the outer medium

In this 1d flat geometry, the hydraulics model with diffusion thus reads:

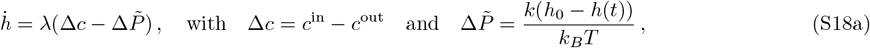

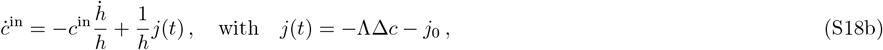

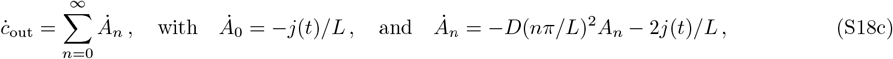

with the initial conditions: *h*(*t* = 0) = *h*_0_, *c*^in^(*t* = 0) = *c*^out^(*t* = 0) = *c*_0_, *A*_*n*_(*t* = 0) = 0.

**FIG. S12.**
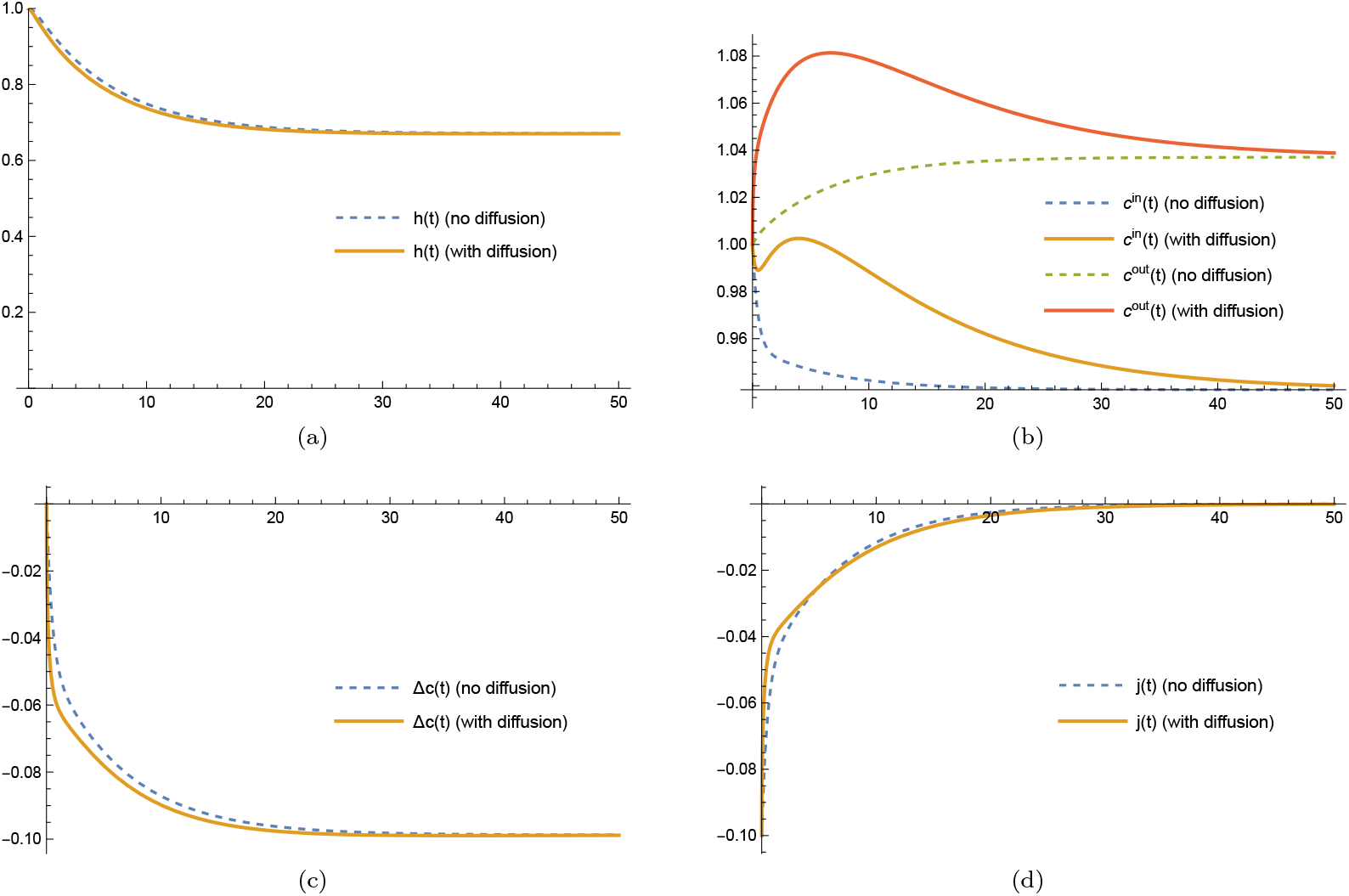
Hydraulic model with (solid lines) and without (dashed lines) diffusion in a 1d flat geometry. (a) Gel thickness *h*(*t*). (b) osmolyte concentration inside *c*^in^(*t*) and outside *c*^out^(*t*) the gel. (c) osmolyte concentration difference Δ*c*(*t*) = *c*^in^(*t*) *− c*^out^(*t*). (d) osmolyte flux in the gel *j*(*t*). Parameters used for the simulations: *λ* = Λ = *k*_*B*_*T* = *D* = *h*_0_ = *c*_0_ = 1, *s* = 0.1, *k* = 0.3, *L* = 10. Number of modes *N* = 100 (converged).

The coupled equations (S18) can be solved numerically by truncating the *A*_*n*_ series at a finite value *N* . Note that truncating the series at the lowest order *N* = 0 reproduces the solution without diffusion. Figure S12 shows an example comparing the dynamics with or without diffusion. Although the concentration inside and ouside have different dynamics (Fig. S12b) with or without diffusion, the gel thickness *h* dynamics (Fig. S12a) and the concentration difference Δ*c* dynamics (Fig. S12c) are not significantly affected.

Importantly, compression is faster with diffusion. Indeed, diffusion in the outer medium increases the osmolyte flux from inside to outside − *j*(*t*), which leads to larger osmotic pressure differences at short time and thus to faster compression.

##### Short-time solution

Finally, to gain some analytical insight, we discuss the short-time solution of the hydraulic model with diffusion. We define *h*(*t*) = *h*_0_ + *δh*(*t*), *c*^in^(*t*) = *c*_0_ + *δc*^in^(*t*) and *c*(*x, t*) = *c*_0_ + *δc*(*x, t*). Introducing the Laplace transform 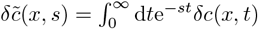, the diffusion equation becomes:

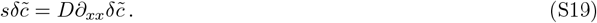

This is an ordinary differential equation on *x* for 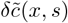, whose solution reads:

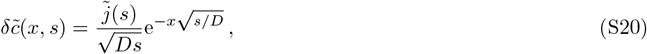

where we have used the boundary conditions 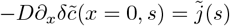 and the simplified boundary condition 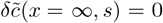, which is reasonable at short time and for large *L*. Then, linearizing the hydraulic equation at first order order in *δc, δh* and using the Laplace transform, we obtain the following system of equations:

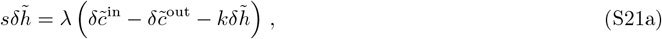

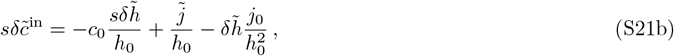

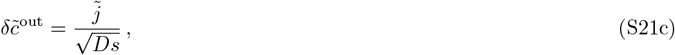

where 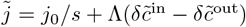. This system can be solved to obtain explicit equations for 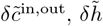 as functions of *s*. The short-time limit of those variables can then be obtained by performing first a series expansion in *s* and then taking the inverse Laplace transform of this series. We thus obtain:

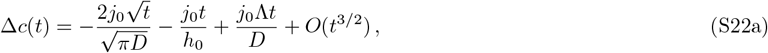

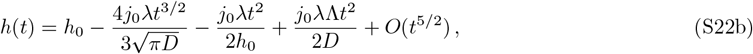

which confirms that compression is faster at short time when diffusion is included.

### II. STOCHASTIC MODEL

#### A. Cell dynamics

In order to account for the strong fluctuations of the normalized spheroid radius around its steady-state observed in the experiments, we considered a stochastic description of division and extrusion of cells at the tissue surface.

We first consider a mean-field description of cells on a surface of fixed size, that undergo divisions *A* → 2*A* with a rate *α* and nonlinear extrusion 2*A* → *A* with rate *β*. A system-size expansion of the master equation leads to the following Itō-Langevin equation for the cell density [7]:

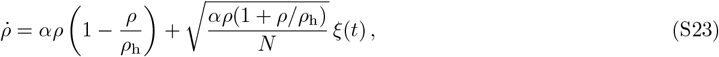

where we have defined a homeostatic density *ρ*_h_ = *α/β*, where *N* is the total cell number and where *ξ*(*t*) is a Gaussian white noise with vanishing mean and variance ⟨*ξ*(*t*)*ξ*(*t*^*′*^)⟩ = *δ*(*t*−*t*^*′*^). Furthermore, we are interested in the fluctuations close to the homeostatic state. To lowest order, we can thus *ρ* ≃ *ρ*_h_, leading to the simplified dynamics:

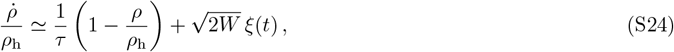

where we have defined *τ* = 1*/α* and *W* = *α/N* . If we now consider that the surface area *A*(*t*) changes with time, the dynamics become:

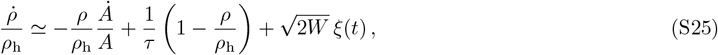

which is the equation given in the main text.

Furthermore, the fluctuating cell density modifies the tissue mechanical properties. We capture this effect at linear order by allowing the tissue surface tension to depend on cell density according to:

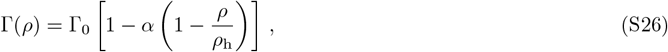

where *α* is a dimensionless parameter and Γ_0_ is the tissue surface tension at the homeostatic density. With our convention, *α >* 0 means that the surface tension is lowered (Γ(*ρ*) *<* Γ_0_) whenever the tissue density is smaller than its homeostatic density (*ρ < ρ*_h_).

The complete model including stochastic cell division thus reads:

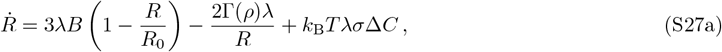

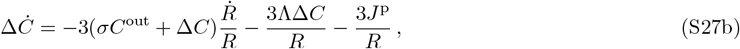

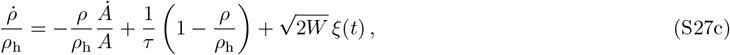

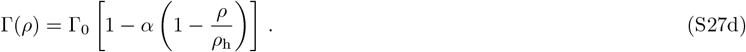

#### B. Alternative feedback

The feedback of cell density fluctuations to the dynamics can also be included as a modification of the tissue permeability to water flows *λ* as:

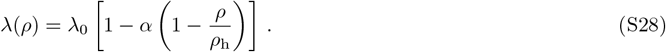

Such feedback, however, only plays a role while the sphere volume is changing, and this feedback thus does not generate radius fluctuations as observed in the experiments. See Sec. III C for details.

We also explored affecting the tissue permeability to ions as:

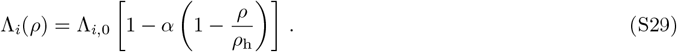

Such feedback leads to radius fluctuations similar to those produced by the tissue surface tension feedback. See Sec. III C for details.

### III. DATA ANALYSIS, FITTING PROCEDURE AND SIMULATIONS

#### A. Steady-state radii

We can analyze the experimental results in the light of the model described in Sec. I. For this purpose, we need to define the initial radius before compression *R*_0_, as well as the steady-state radius after compression *R*^*\**^:

- We define the initial radius before compression *R*_0_ as the time average of the radius before the beginning of the compression stage. The first time point of the compression stage is defined as the first point which is followed by 7 successive time points where the radius decreases.
- We considered two different definitions of the steady-state radius *R*^*\**^ after the compression stage. A simple choice is to define it as *R*^*\**^ = *R*_min_ where *R*_min_ is the minimal radius after the compression stage. Another possibility is to define *R*^*\**^ = *R*_mean_ where *R*_mean_ is the mean value after the end of the compression stage, where the end of the compression stage is defined as the time point at which the radius as decrease to the 3/4 of *R*_min_.

An example of the definition of these quantities for an osmotic shock experiment is displayed in Fig. S13.

Using Eq. (S6), we fitted the experimental results with two fitting parameters *J* and *γ*. We obtain *J*≃ 3.3 10^*−*4^ mM^*−*1^ and *γ* ≃ 3.8 10^*−*2^ for the case *R*^*\**^ = *R*_min_ and *J* ≃ 3.0 10^*−*4^ mM^*−*1^ and *γ* ≃ 3.2 10^*−*2^ for the case *R*^*\**^ = *R*_mean_. To be consistent with the fact that the tissue dynamics is stochastic, we present the case *R*^*\**^ = *R*_mean_ in the main text.

#### B. Fitting of the dynamics

The dynamics predicted by our model can then be compared to the experimental data. For this purpose, we solved the dimensionless equation (S8) using the NDSolve function provided by the software Mathematica with the initial conditions *r*(*t*_i_) = *r*_i_ and *δc*(*t*_i_) = 0. In the simulation, an osmotic shock was described by solving Eq. (S8) with 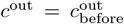 up to from the initial time *t*_*i*_ to the osmotic shock time 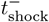. Then a new simulation with 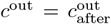 was started at time 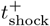, with initial radius 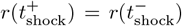 and initial concentration difference 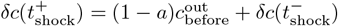 where 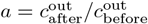.

**FIG. S13.**
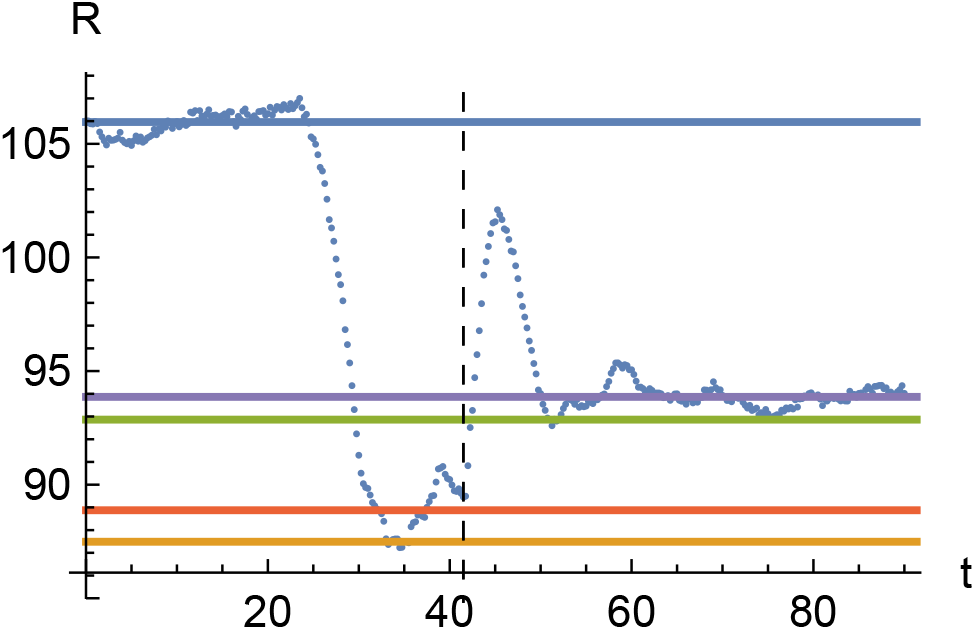
Example of a osmotic shock experiment (blue dots). The black dashed vertical line indicates the time point at which the osmotic shock has been applied. The solid lines show the different definitions used in the text. Blue line: average radius before the shock, *R*_0_. Orange line : minimum radius before shock,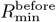. Red line : average radius before shock, 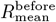. Green line : minimum radius after shock, 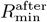. Red line : average radius after shock, 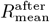.

**FIG. S14.**
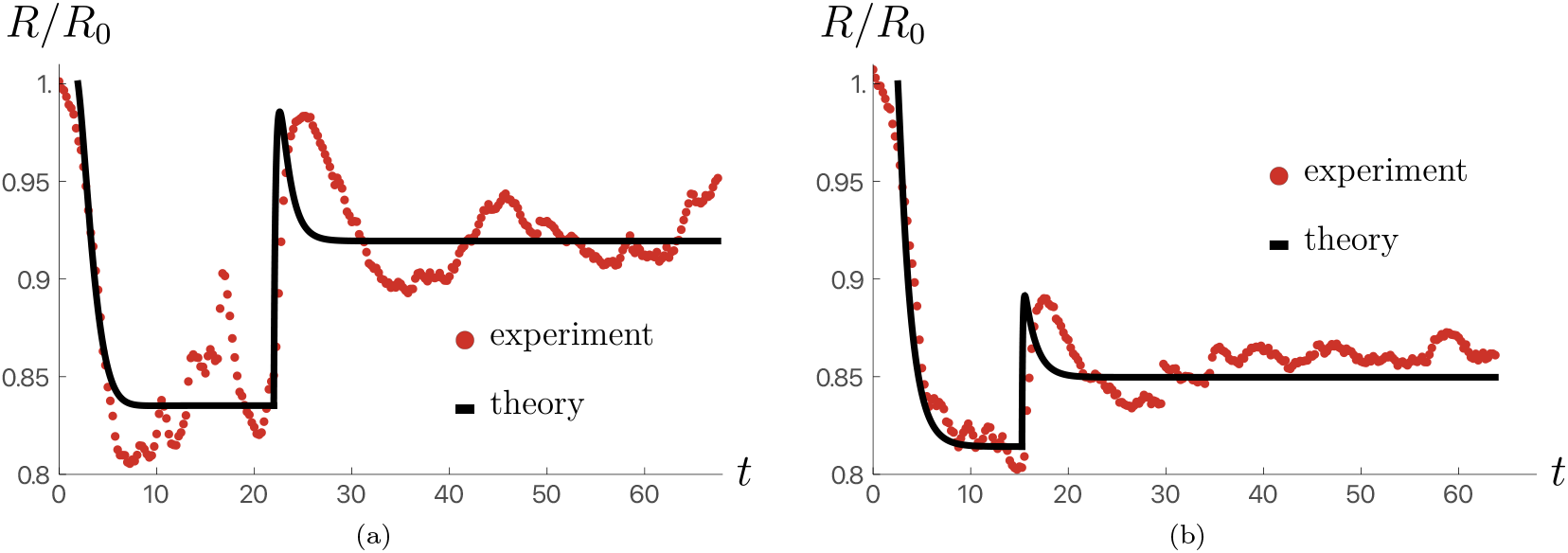
Radius dynamics predicted by Eq. (S8) (solid black line) and comparison with osmotic shock experiments (red dots). Osmotic shock from 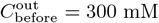 to 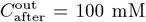. Parameters used for the simulations: *γ* = 0.0186, *κ* = 0.6, *σ* = 0.04, *j* = 1.9, 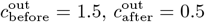. **(b)** Osmotic shock from 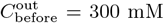 to 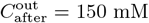. Parameters used for the simulations: *γ* = 0.051, *κ* = 3.52, *σ* = 0.059, *j* = 0.45, 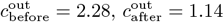.

From the steady-state radii fit discussed above, the average over several experiments of the dimensionless parameter *γ*, as well as the product *jc*^out^*σ* = *JC*^out^ were determined. For fitting indiviual dynamics as discussed below, we nonetheless allow *γ* and *jc*^out^*σ* to deviate from these values.

Figure S14 shows two examples of experimental data and their corresponding fits. We note that a family of similarly good fits can be obtained by varying both *σ* and *κ*, provided *σ* ≲ 0.5.

#### C. Stochastic simulations

**FIG. S15.**
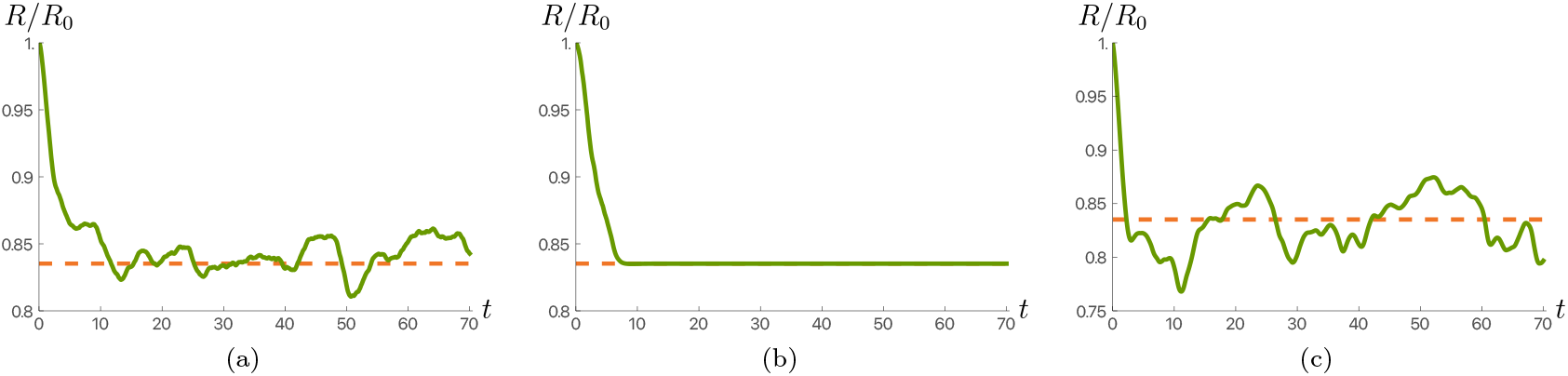
Example of simulation of the stochastic model (S27). Solid green line: normalized radius; orange dashed line: steady-state radius in the absence of stochasticity. **(a)** Stochastic model with tissue tension feedback. **(b)** Stochastic model with tissue permeability to water flow feedback [Eq. (S27d) replaced by Eq. (S28)]. **(c)** Stochastic model with tissue permeability to ion feedback [Eq. (S27d) replaced by Eq. (S29)]. Parameters used for the simulations: *κ* = 0.6, *σ* = 0.04, *γ* = 0.0186, *j* = 1.9, *c*^out^ = 1.5, *α* = 2, *τ* = 5; *W* = 0.01.

The stochastic simulations of Eq. (S27) were performed using the ItoProcess and RandomFunction functions of the software Mathematica. Including the multiplicative noise of Eq. (S23) or using the simplified additive noise as appearing in Eq. (S24) yields qualitatively similar results, and we therefore used the additive noise.

Figure S15 shows examples of realization of the stochastic equation (S27) with different feedback from cell density to the dynamics. Figure S15a shows an example where the feedback stems from the tissue surface tension (S26), as used in the main text. Figure S15b shows an example where the feedback stems from permeability to water flow [Eq. (S27d) replaced by Eq. (S28)]. Such feedback only plays a role while the sphere volume is changing, and it thus does not generate radius fluctuations as observed in the experiments. Figure S15c shows an example where the feedback stems from tissue permeability to ions [Eq. (S27d) replaced by Eq. (S29)]. This type of feedback also leads to radius fluctuations similar to those observed in the experiments.

Finally, to obtain a stochastic simulation displayed in the main text and that resemble the experimental osmotic shock experiment, we ran *N* = 1000 realizations from time *t* = *t*_i_ to *t* = *t*_shock_ and picked the one minimizing a quadratic cost function that penalizes the distance to the experimental points^1^. We then ran *N* = 1000 realizations for the osmotic shock part from *t* = *t*_shock_ to *t* = *t*_end_ and again picked the one minimizing the cost function.

#### D. Parameters used in the figures

We summarize here the parameters used to produce Fig. 5 of the main text:

- Fig. 5B: Obtained using Equations (S11) and the eigenvalues from the stability matrix (S12). Parameters used are: *κ* = 0.6, *σ* = 0.04, *c*^out^ = 1.5.
- Fig. 5E: Radius dynamics obtained from Eq. (S8). Parameters used: *γ* = 0.0186, *κ* = 0.6, *σ* = 0.04, *j* = 1.9, 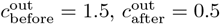.
- Fig. 5F: Stochastic simulation with tissue tension feedback, Eq. (S27). Parameters used for the simulations: *κ* = 0.6, *σ* = 0.04, *γ* = 0.0186, *j* = 1.9, 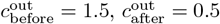, *α* = 2, *τ* = 5; *W* = 0.01.

## Footnotes

1 The cost function *C* reads *C* = _*n*_ *r*^exp^(*t*_*n*_) *− r*^simu^(*t*_*n*_) ^2^ where *r*(*t*) = *R*(*t*)*/R*_0_ and the *t*_*n*_ are the experimental time points.

